# The immune response to lumpy skin disease virus in cattle is influenced by inoculation route

**DOI:** 10.1101/2022.09.22.509006

**Authors:** Petra C. Fay, Najith Wijesiriwardana, Henry Munyanduki, Beatriz Sanz-Bernardo, Isabel Lewis, Ismar R. Haga, Katy Moffat, Arnoud H. M. van Vliet, Jayne Hope, Simon Graham, Philippa M. Beard

**Affiliations:** The Pirbright Institute, Ash Road, Pirbright, GU24 0NF, UK; School of Veterinary Medicine, University of Surrey, Daphne Jackson Road, Guildford, GU2 7AL, UK; The Roslin Institute, Easter Bush, University of Edinburgh, EH25 9RG, UK

**Keywords:** Virus, poxvirus, bovine immunity, lumpy skin disease, LSDV, neutralising antibodies, humoral immunity, cell-mediated immunity

## Abstract

Lumpy skin disease virus (LSDV) causes severe disease in cattle and water buffalo and is transmitted by hematophagous arthropod vectors. Detailed information of the adaptive and innate immune response to LSDV is limited, hampering the development of tools to control the disease. This study provides an in-depth analysis of the immune responses of calves experimentally inoculated with LSDV via either needle-inoculation or arthropod-inoculation using virus-positive *Stomoxys calcitrans* and *Aedes aegypti* vectors. Seven out of seventeen needle-inoculated calves (41%) developed clinical disease characterised by multifocal necrotic cutaneous nodules. In comparison 8/10 (80%) of the arthropod-inoculated calves developed clinical disease. A variable LSDV-specific IFN-γ immune response was detected in the needle-inoculated calves from 5 days post inoculation (dpi) onwards, with no difference between clinical calves (developed cutaneous lesions) and nonclinical calves (did not develop cutaneous lesions). In contrast a robust and uniform cell-mediated immune response was detected in all eight clinical arthropod-inoculated calves, with little response detected in the two nonclinical arthropod-inoculated calves. Neutralising antibodies against LSDV were detected in all inoculated cattle from 5-7 dpi. Comparison of the production of anti-LSDV IgM and IgG antibodies revealed no difference between clinical and nonclinical needle-inoculated calves, however a strong IgM response was evident in the nonclinical arthropod-inoculated calves but absent in the clinical arthropod-inoculated calves. This suggests that early IgM production is a correlate of protection in LSD. This study presents the first evidence of differences in the immune response between clinical and nonclinical cattle and highlights the importance of using a relevant transmission model when studying LSD.

## 1 Introduction

Lumpy skin disease (LSD) is a high consequence disease of cattle and water buffalo caused by lumpy skin disease virus (LSDV), a large double-stranded DNA poxvirus within the *Capripoxvirus* genus (1, 2). Since it was first described in southern Africa in 1929 LSDV has progressively spread throughout Africa where it is now endemic, and more recently into the Middle East, Russia, and Europe (3, 4). Since 2019 LSDV has spread throughout Asia establishing itself as a major emerging transboundary pathogen (5–9).

LSD is characterised by multifocal necrotic cutaneous lesions accompanied by lymphadenopathy, lethargy, pyrexia, weight loss and reduced milk production (10–12). The morbidity and mortality rates in outbreaks of LSD vary from approximately 10-20% morbidity and 1-10% mortality (13–15). The pathology is focused on the skin as a multifocal dermatitis with vasculitis of dermal blood vessels, resulting in full-thickness necrosis of the dermis and epidermis (11). Unusually for a poxvirus, direct transmission of LSDV is rarely documented however LSDV can be transmitted by hematophagous arthropods including *Aedes aegypti* mosquitoes, *Rhipicephalus microplus* ticks and *Stomoxys calcitrans* stable flies (14, 16–20). The main methods for the control and prevention of LSDV are effective surveillance programmes to detect outbreaks, widespread use of live attenuated vaccines, and ‘stamping out’ of infected herds (20–22). LSD has significant socioeconomic implications, particularly in low- and middle-income countries, due to the drop in milk yield, reduced meat production, poor hide quality, cost of control measures and trade restrictions (2, 11, 23). Efforts to develop better tools for the detection, control and prevention of LSD are hampered by a poor understanding of the immune response to LSDV (24, 25).

Characterisation of the adaptive immune response to LSDV is limited. The three species of capripoxviruses (LSDV, goatpox virus and sheeppox virus) are genetically very similar and serologically indistinct. Previous studies examining immune responses to capripoxvirus vaccination indicate both a cell-mediated and humoral immune response is generated (26–32). Strong antibody responses are produced by cattle that are vaccinated with live attenuated strains of LSDV. Both binding antibodies (bAbs) and neutralising antibodies (nAbs) are detected and titres can vary markedly between clinical and nonclinical animals (26, 31, 33–35). Neutralising antibodies play a role in long-term protection post-vaccination, similar to other poxviruses (36, 37), and have been shown to be long-lasting in follow-up studies in goats and sheep vaccinated against sheeppox virus (28, 30, 31).

The role of cell-mediated immunity (CMI) in LSD is particularly poorly understood (2, 38, 39). Primarily driven by T lymphocytes, this immune response results in the production of key cytokines including type II IFN (IFN-γ) which is produced by CD4+ helper T cells, CD8+ cytotoxic T cells, ɣδ T cells, natural killer T cells, and NK cells (40). IFN-γ and other cytokines induced by the CMI response have a range of functions including the activation of NK cells and macrophages and inducing the class switching of immunoglobulins from activated plasma B cells (41–44).

Previous literature reports the detection of a CMI response following inoculation with wildtype or attenuated strains of LSDV. The first evidence of a CMI response to LSDV was reported in 1995 with description of a delayed-type hypersensitivity (DTH) reaction following virus inoculation (45). Recently more complex immune assays have quantified IFN-γ, a key biomarker of the CMI response, in cattle that had been vaccinated or challenged, or both (46–48), with evidence to suggest the involvement of CD4^+^ and CD8^+^ T cells in the production of IFN-γ (46). However, the kinetics and magnitude of CMI response against LSDV, and the role it plays in protection against disease, is not yet understood.

Following experimental infection with LSDV, a proportion of cattle develop clinical disease characterised by cutaneous lesions whilst other animals remain nonclinical (11, 18). It remains unknown what mechanisms drive these differences and if the elicited immune response to infection plays a role in the presentation of clinical disease. In this study, we have evaluated the humoral and CMI responses in cattle experimentally inoculated by two different routes, either by needle inoculation as described previously (11, 19) or by LSDV-positive blood-feeding arthropods (*S. calcitrans* and *Ae. aegypti*), a route which is more representative of virus transmission in the field (18). In-depth data was obtained that characterises the adaptive immune response to LSDV, correlates differences with clinical outcome, and provides critical insight into the progression of disease in the host. The information obtained provides highly relevant detail that can be applied to developing improved disease control measures for an increasingly important transboundary disease.

## 2 Materials and Methods

### Ethical Statement

This work was conducted under license P2137C5BC from the UK Home Office at The Pirbright Institute according to the Animals (Scientific Procedures) Act 1986, and approved by The Pirbright Institute Animal Welfare and Ethical Review Board.

### Viruses

The LSDV strain used for inoculation of all animals was sourced from the WOAH Capripoxvirus Reference Laboratory at The Pirbright Institute and originated from a LSD outbreak in eastern Europe in 2016. The Neethling (35) and Cameroon strain of LSDV were sourced from the WOAH Capripoxvirus Reference Laboratory at The Pirbright Institute. Viruses were grown and titred on MDBK cells as described previously (49). Mock virus preparations were produced in an identical manner from uninfected MDBK cells for use as a negative control in immunological assays

### Experimental infection of calves with LSDV

Male castrated Holstein-Friesian calves were included in the study. Median age and weight in group A were 104 days old and 145 kg, in group B 124 days old and 176 kg, in group C 140 days and 157 kg, in group D 97 days and 126 kg, in group RA and RS 96 days and 120 kg (Table 1). The animals were sourced from a commercial high health herd and confirmed as negative for BVDV via PCR prior to study commencement. The animals were housed in the high-containment animal facility (SAPO4) at The Pirbright Institute. Up to five calves were housed in one room (22m^2^) with appropriate bedding material (Mayo Horse Comfort) and environmental enrichment, such as rubber toys and hollow containers filled with hay, provided. Light/dark cycle was 12:12 h, temperature was held between 10°C to 24°C, and humidity 40% to 70%. Animals were fed concentrated rations twice daily and given *ad lib* access to hay and water.

**Table 1.** Cattle study experimental design.

| Cattle Study Groups and Animal Numbers |  |  |  |  |  |  |
| --- | --- | --- | --- | --- | --- | --- |
|  | A | B | C | D<br>'Donors' | RA<br>'Recipients' | RS<br>'Recipients' |
| **Uninfected** | A1 | B6 | C11 | none | none | none |
| **Inoculated** | A2, A3, A4,<br>A5 | B7, B8, B9,<br>B10 | C12, C13,<br>C14, C15 | D1, D2,<br>D3, D4, D5 | RA1, RA2,<br>RA3, RA4,<br>RA5 | RS1, RS2,<br>RS3, RS4,<br>RS5 |
| **Inoculation**<br>**Route** | Intravenous<br>and intradermal | Intravenous<br>and<br>intradermal | Intravenous<br>and<br>intradermal | Intravenous<br>and<br>intradermal | *Stomoxys*<br>*calcitrans* | *Aedes aegypti* |

\*Study duration for the intravenous and intradermal cattle groups was 21 days and recipients 29 days.

Four experimental studies are reported in this manuscript. The design, clinical outcomes, and pathology from the first three studies (A-C) have been reported previously (11, 18, 50). Briefly, in each of these studies a group of 5 calves were randomly assigned to treatment (n=4) or non-treatment (n=1) groups. The 4 treated calves were inoculated by needle injection with 3 mL of LSDV at a concentration of 1 × 10^6^ PFU/mL. Two mL (2 × 10^6^ PFU) of virus was inoculated intravenously (IV) into the jugular vein, and 1 mL (1 × 10^6^ PFU) of virus injected intradermally (ID) into 2 sites on each side of the neck (0.25 mL in each site). The untreated animal in each experiment was not inoculated. In all three experiments blood-feeding arthropods were fed on the skin of some of the inoculated calves as described previously (11, 18, 50).

In the fourth experimental study fifteen calves were assigned into three groups of 5 (groups D, RA and RS). The 5 calves in group D were inoculated as described above with 3 × 10^6^ PFU/mL intravenously and intradermally. One calf in group D (D5) developed over 100 cutaneous lesions and was then used as a “donor” animal. *Stomoxys calcitrans* and *Aedes aegypti* insects were bred at The Pirbright Institute as described previously (18, 50). Adult insects were fed on the cutaneous lesions of calf D5 for a maximum of 40 sec before being interrupted and within a maximum of 1 h fed on the calves in group RA (n=5) and RS (n=5), as described in Table 1.

### Peripheral blood mononuclear cell isolation

Heparinised blood was diluted with PBS at a ratio of 1:1 and overlaid onto Histopaque 1083 (Sigma-Aldrich, Merck) density gradient medium using SepMate™-50 centrifugation tubes (Stem Cell Technologies). Samples were centrifuged at 1500 × *g* for 30 min at 20°C with the brake off. Peripheral blood mononuclear cells (PBMCs) were aspirated from the interface and washed twice with PBS centrifuging at 1000 × *g* for 10 min at 20°C. After the final wash, PBMCs for T-cell ELISpot analysis were resuspended in 3mL in RPMI 1640 medium GlutaMAX™ (ThermoFisher Scientific) only and those for B-cell ELISpot analysis were resuspended in 3mL RPMI 1640 medium GlutaMAX™ (ThermoFisher Scientific) supplemented with 10% heat-inactivated horse serum (Gibco™) and 100IU/mL penicillin and 100µg/mL streptomycin (ThermoFisher Scientific). Viable cells were counted using a Cellometer cell counter (Nexcelom Bioscience).

### Short-wave UV inactivation of LSDV

Live LSDV Neethling was placed in a 6-well plate on ice. A hand-held UV lamp emitting shortwaves (245 nm) was placed approximately 4 inches above the lidless 6-well plate and the virus irradiated for 5 minutes. Any residual viral infectivity was determined by plaque assay.

### Bovine Type I IFN Mx/CAT reporter assay

To determine the levels of biologically active bovine type I IFN, a Mx/chloramphenicol acetyltransferase (Mx/CAT) reporter assay was used as described previously (51). MDBK-t2 cells (gifted by Veronica Carr, The Pirbright Institute) were cultured in MEM (Sigma) containing 10% FBS, 100 IU/mL penicillin, 100µg/mL streptomycin, and 10 µg/mL of blasticidin (InvivoGen). MDBK-t2 cells were seeded into 24-well plates at 1 × 10^5^ / well and incubated at 37°C in a 5% CO_2_ incubator overnight. Serum samples (500 µL) were heat-inactivated at 56°C for 30 mins using a heat block. Heat-inactivated sera samples (250 µL / sample) were incubated on MDBK-t2 cells in media and recombinant bovine IFN-α standards (15.6-500 IU/mL;) were added to MDBK-t2 cells at a 1:1 ratio to form a standard curve. MDBK-t2 cells were incubated with the sera and recombinant bovine IFNα standards overnight at 37°C in a 5% CO_2_ incubator. CAT expression was determined using an ELISA kit in accordance with the manufacturer’s instructions (Roche). Bovine IFN-α standards were used to construct a type I IFN standard curve to interpolate sera sample IFN levels.

### PBMC Interferon Gamma Release Assay (IGRA)

PBMCs were seeded at 5 × 10^5^ cells per well in 96-well round bottom plates and stimulated with either the positive controls (pokeweed mitogen (PWM) or concanavalin A (Con A)), negative controls (cell culture media or mock infected cells) or SW-UV inactivated LSDV (diluted 1:10). These cells were incubated for 24 h at 37°C in a 5% CO_2_ incubator after which the PBMCs were centrifuged at 300 × *g* for 5 min and the supernatants were harvested and stored at −80°C.

ELISA plates were coated with 50 µL/well of 5 µg/mL of mouse anti-bovine IFN-γ monoclonal antibody (mAb) (CC330, Bio-Rad Antibodies) by overnight incubation at 4°C. Plates were washed five times with washing buffer (PBS and 0.05% Tween20 (Thermo Fisher Scientific). 100 µL/well of blocking buffer (PBS, 0.05% Tween-20, 0.5% BSA) was added and plates incubated for 1 hour at 37°C. Blocking buffer was removed and 100 µL/well of samples and IFN-γ standards (recombinant bovine IFN-γ; Bio-Rad Antibodies) starting at 300 ng/mL followed by two-fold dilutions added. Samples were diluted at 1:2 in carrier buffer (PBS, 0.05% Tween 20, 0.5% BSA). Plates were incubated for 1 hour at 37°C and then washed five times with washing buffer. Biotinylated mouse anti-bovine IFN-γ mAb (CC302; Bio-Rad Antibodies) diluted to 0.5µg/mL in carrier buffer was added (100 µL/well) and plates incubated for 1 hour at 37°C. After five washes with washing buffer, streptavidin-HRP conjugate (N-100, Thermo Fisher Scientific) diluted 1:1500 in carrier buffer was added (50 µL/well) and incubated for 30 min at 37°C. After five final washes, 50 µL/well of 3,3’,5,5’-tetramethylbenzidine (TMB) substrate (Thermo Fisher Scientific) was added and incubated at RT for 5 min. Substrate development was followed by the addition of stop solution (1N sulfuric acid). Optical densities were immediately read in a spectrophotometer at 450 nm.

### Whole Blood Interferon Gamma Release Assay (IGRA)

Heparinised whole blood was collected at selected timepoints and stimulated with 20 µL of live LSDV Neethling (4 × 10^7^ PFU/mL) overnight at 37°C in a 5% CO_2_ incubator. PWM was used a positive control and PBS as a negative control. The following day, plasma was collected and used to test for secretory IFN-γ using the ID Screen® Ruminant IFN-g ELISA following the manufacturer’s guidelines (Innovative Diagnostics).

### Quantification of LSDV in blood

Whole blood (EDTA) was collected to investigate viremia. Blood was collected 3 days before needle inoculation, on day 5 post-infection (dpi), and every second day until day 21 dpi. For the insect inoculated recipient calves, blood was taken a day before insect feeding and every second day until day 28 dpi. Viral DNA was extracted from blood samples using the KingFisher Flex extraction instrument (Thermo Fisher Scientific) and MagMax Core Extraction kit (A32700; Thermo Fisher Scientific) using Workflow A for DNA extraction from whole blood. The MagMAX_CORE_No_Heat protocol was used as per the manufacturer’s guidelines, with minimal modifications. DNA was eluted in 50 µL of elution buffer. Plasmids containing LSDV ORF068 (19AEL6VP; Thermo Fisher Scientific) and bovine cytochrome B (19AEL7QP; Thermo Fisher Scientific) were linearised and used to generate standards. To determine viral load in the skin, 2mm punch biopsies from recipient calves were collected on each day of insect feeding on donor calves and at days 9 and 15 days post infection on insect inoculated calves. The biopsies were digested by adding 90 µL of proteinase K buffer (4489111, Thermo Fisher Scientific) and 10 µL proteinase K (25530049, Thermo Fisher Scientific) to the skin biopsy. The sample was incubated for 2 hours at 55°C. Tubes were lightly vortexed during incubation and centrifuged briefly. 20 µL of MagMAX Core magnetic beads (Thermo Fisher Scientific) were added to the lysate and mixed. 100 µL of the bead and lysate mixture was transferred to the KingFisher deep well plates (Thermo Fisher Scientific) containing 350 µL of lysis solution and 350 µL of binding solution. Extraction was done as outlined in the MagMAX core extraction kit and the MagMAX_CORE_No_Heat protocol was used to extract DNA from the samples. DNA was eluted in 50 µL of elution buffer. A TaqMan multiplex PCR assay designed to amplify LSDV068 using forward and reverse primers 5’ GGCGATGTCCATTCCCTG 3’ and 5’ AGCATTTCATTTCCGTGAGGA 3’ respectively and a probe - ABY 5’ CAA TGG GTA AAA GAT TTC TA3’ QSY was used to detect LSDV. A forward primer – GTAGACAAAGCAACCCTTAC and a reverse primer -GGAGGAATAGTAGGTGGAC for bovine Cytochrome B and a probe - FAM 5’TTA TCA TCA TAG CAA TTG CC 3’ MGBNQF was used for the detection of bovine cytochrome B. TaqMan Multiplex master mix (4461884, Thermo Fisher Scientific) was used with MUSTANG PURPLE as the passive reference dye. In brief, the reaction setup was as follows; 5 µL of template was used in a total of 20 µL reaction. Final primer concentrations were 500 nM and 250nM for probes. An initial denaturation at 95°C for 20 seconds was carried out and 50 cycles of denaturation at 95°C for 15 seconds, annealing and signal acquisition at 58°C for 1 min were done. Viremia was expressed as genome copies per/mL using the equation below.

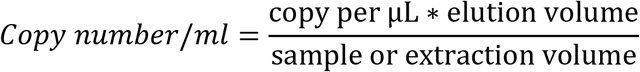

A lower detection limit of 500 copies/mL was set. Viremia was analysed in duplicate and results presented as mean copies per/mL.

### Flow cytometric analysis of LSDV-specific T-cell responses

PBMCs were seeded at a density of 5 × 10^5^ cells/well in 96-well round-bottom tissue culture plates. Each treatment condition i.e., positive control (PMA/Ionomycin), negative control (media), LSDV Neethling (live virus), or UV-inactivated LSDV Neethling, was tested in triplicate wells. Once the cells were seeded, wells were stimulated with LSDV Neethling (live virus) at MOI =1, or an equivalent dose of UV-inactivated LSDV Neethling virus followed by overnight incubation at 37°C in a 5 % CO2 incubator. After the addition of 10 μg/mL Brefeldin A (BFA) (Merck) to all wells, and 10 μg/mL phorbol 12-myristate 13-acetate (PMA) and Ionomycin (Merck) to corresponding positive control wells, cells were incubated for a further 4 hours at 37°C in a 5 % CO_2_ incubator. After centrifugation at 300 × g for 5 min and removal of the supernatant, cells were washed once in FACS buffer (PBS and 1% BSA) and then surface labelled by incubation with anti-bovine CD8β-RPE mAb [CC58; Bio-Rad Antibodies), anti-bovine CD4: FITC mAb [CC8; Bio-Rad Antibodies] diluted in FACS buffer for 10 min at RT in the dark. Dead cell staining was performed using a LIVE/DEAD™ Fixable Violet Dead Cell Stain Kit following manufacturer guidelines (Thermo Fisher Scientific). Cells were then washed twice in FACS buffer and then fixed in BD CytofixCytoPerm (BD Biosciences) for 20 min at RT in the dark. Cells were then washed twice in BD PermWash solution (BD Biosciences). Permeabilised cells were immediately labelled by incubation with anti-bovine IFN-γ-APC mAb (CC302; Bio-Rad Antibodies) diluted in BD PermWash solution, for 30 min at 4°C in the dark. After washing twice in BD PermWash solution, cells were resuspended in PBS and immediately analysed. Data was collected using the MACSQuant10 Analyzer flow cytometer (Miltenyi Biotec) equipped with a 405nm (violet), 488nm (blue) and 640nm (red) laser. Data was analysed with FCSexpress7 software (DeNovo) and samples were gated on cells (SSC-A vs FSC-A), singlets (SSC-A vs SSC-H) and then CD4^+^ cells were determined as FITC positive (blue: 525/50), CD8^+^ cells were determined as RPE positive (blue: 585/40) and IFN-γ positive cells were determined as APC positive (red: 655-730; Supplementary Figure 1).

### IFN-γ ELISpot assay

Multiscreen-IP 0.45 µM multiwell filter plates (#MAIPS4510, Merck) were coated with 50 µL/well of 2 µg/mL of mouse anti-bovine IFN-γ mAb (CC330 Bio-Rad Antibodies) diluted in carbonate coating buffer (pH 9.6) for 2 hours at RT. Plates were washed 5 × with 200 µL PBS/well and then blocked with 50 µL/well of 4% semi-skimmed powdered milk (Marvel) in PBS for 2 hours at RT. Plates were washed 5 times with 200 µL PBS, 100 µL of PBS/well added, plates sealed and stored at 4°C overnight. PBMCs were resuspended to a concentration of 5 × 10^5^ cells/mL in RPMI 1640 medium GlutaMAX™ (Thermo F isher Scientific). In a separate plate, two-fold dilutions of the cells were prepared from this at 1:2 and 1:4 and 50 µL added to corresponding wells of the coated multiscreen plate. Negative control wells containing PBS only and positive control wells containing 2 µ g/mL of PWM; Sigma-Aldrich) were included for each timepoint. Plates were incubated at 37°C in a 5% CO2 incubator overnight. The following day plates were washed 5 times with 200 µL/well of PBS containing 0.05 % Tween20 (Thermo Fisher Scientific). To each well 50 µL of 2 µg/mL of biotin-conjugated mouse anti-bovine IFN-γ mAb (CC302, Bio-Rad Antibodies) diluted in PBS was added and plates incubated for 2 hours at RT. Plates were then washed as previously and 50 µL/well of streptavidin AP (Southern Biotech) diluted 1:1000 in PBS was added and plates incubated at RT for 1 hour. Plates were washed as previously. To each well 50 µL/well of alkaline phosphatase substrate (Bio-Rad) was added and incubated at RT for approximately 15 min or until spots developed. The reaction was stopped by immersing plates in tap water. Plates were left to dry overnight at RT, face down before enumeration of spots using the ImmunoSpot 7.0 reader and ImmunoSpot SC suite (Cellular Technology Limited). Results were expressed as the mean number of spot-forming cells/million PBMC.

### LSDV antibody ELISA

LSDV-specific antibodies were detected using the commercial ELISA kit ID Screen^®^ Capripox Double Antigen Multi-species ELISA kit following manufacturer instructions (Innovative Diagnostics).

### LSDV Fluorescent Virus Neutralisation Test (FVNT)

To measure nAb titres a FVNT was used to test serum samples collected at each time point following the protocol described by previously (31). nAb titres were determined as the highest reciprocal dilution at which no foci were identified, indicative of complete neutralisation. Partial neutralisation was determined by counting the number of fluorescent foci at each dilution using a cut-off of 50 foci per well and converting this into a neutralisation percentage.

### Antibody secreting cell (ASC) B cell ELISpot

Multiscreen-IP 0.45 µM multiwell filter plates (#MAIPS4510, Merch) were activated with 50 µL/well of 35% ethanol for 30 seconds. Plates were washed 5 times with 200 µL/well sterile water. Plates were coated with 50 µL/well of LSDV Cameroon (4 × 10^8^ PFU/mL) diluted 1:120 in 0.1 M carbonate bicarbonate buffer (pH 9.6) and incubated at 4°C overnight. The following day, plates were washed 5 times with 200 µL/well with PBS and blocked with 100 µL/well of PBS containing 4% powdered skimmed milk (Marvel) and incubated at RT for 1 hour. Plates were washed as previously. Isolated PBMCs were resuspended in RPMI 1640 medium GlutaMAX™ supplemented with 10% heat-inactivated horse serum and 100IU/mL penicillin and 100µg/mL streptomycin (all supplied by Thermo Fisher Scientific) to a density of 5 × 10^5^ cells/mL. In a separate round bottom 96-well tissue culture plate, two-fold dilutions of the cells were prepared from this at 1:2 and 1:4 and 50 µL added to corresponding wells of the coated multiscreen plate including a negative control of uninfected cell lysate and incubated at 37°C in a 5 % CO_2_ incubator overnight. Plates were washed with 200 µL/well of PBS containing 0.05% Tween-20. To corresponding wells, 100 µL/well of biotinylated goat anti-bovine IgG-heavy and light chain antibody or sheep anti-bovine IgM antibody (Bethyl Laboratories) diluted 1:1500 in PBS was added and plates incubated at RT for 2.5 hours. Plates were washed as previously and 100 µL/well of streptavidin-AP (Southern Biotech) diluted 1:1000 was added, and plates incubated at RT for 1 hour. Plates were washed as previously and to each well 50 µL/well of colorimetric alkaline phosphatase (AP) substrate (BioRad) was added followed by incubation at RT for approximately 15 min or until spots first develop. The reaction was stopped by immersing plates in tap water. Plates were left to dry overnight at RT, face down before enumeration of spots using the ImmunoSpot 7.0 reader and ImmunoSpot SC suite (Cellular Technology Limited). Results were manually validated for false-positive results and expressed as the mean number of ASCs/million.

### Statistical Analysis

Statistical significance for differences was performed using a two-way analysis of variance (ANOVA) followed by Sidak’s multiple comparison test using GraphPad Prism software version 8.2.0 for Windows.

## 3 Results

### A combined intravenous and intradermal challenge with LSDV results in clinical and nonclinical disease in calves

A total of 17 calves in groups A-D were needle inoculated with LSDV via both intravenous and intradermal routes and then monitored for clinical disease. In total, 7 calves (41%) developed clinical disease. Clinical findings for groups A - C have been described previously (11, 18, 50). The clinical outcomes in group D are summarised in Figure 1 and were consistent with those seen previously in groups A - C. Three of the 5 calves in group D developed clinical disease (D2, D4 and D5), defined as cutaneous lesions distant from the inoculation site. These lesions were first detected on calves D2 and D4 at 5 days post-infection (dpi) and on D5 at 6 dpi (Figure 1A). The two remaining calves (D1 and D3) did not develop clinical disease and were classified as nonclinical (Figure 1B). Calf D5 developed the most severe disease with over 100 cutaneous lesions present from 13 dpi until the end of the study period (21 dpi). A rise in rectal temperature was observed from 3dpi in the clinical calves and remained elevated until 10, 13 or 16 dpi in D2, D4 and D5 respectively (Figure 1C). No increase in rectal temperature was detected in the nonclinical calf D1, however D3 exhibited a temperature spike of 39.8°C at 4 dpi (Figure 1D). A brief period of pyrexia in nonclinical animals has been noted in previous studies by ourselves and others (19).

**Figure 1.**
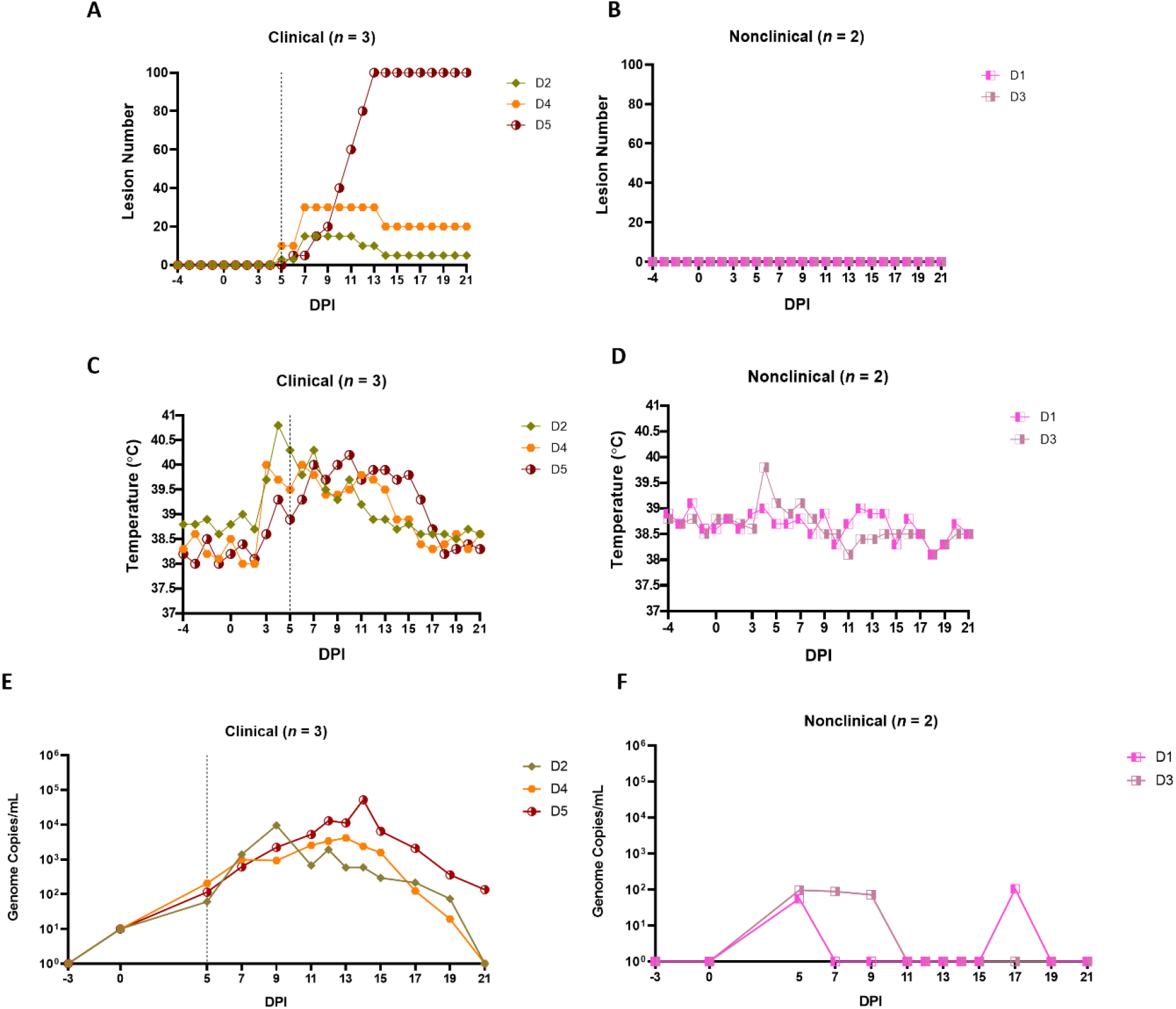
Clinical and virological outcomes of clinical and nonclinical calves after needle-inoculation with LSDV. The number of cutaneous lesions (**A** and **B**) were recorded each day, up to a maximum of 100 lesions. The rectal temperature of clinical and nonclinical calves (**C** and **D**) was recorded daily. The number of LSDV genome copies in the blood of each calf was quantified by qPCR. A vertical dotted line represents the first day cutaneous lesions were noted in the clinical calves.

Lymphadenopathy of the prescapular lymph nodes was noted in all inoculated calves from 2-3 dpi onwards and was more pronounced in the clinical calves. Lymphadenopathy of both left and right prefemoral lymph nodes was also noted in calf D5 from 11 dpi. Calf D5 exhibited lethargy on 12-16 dpi and loss of body condition from around 10 dpi onwards. The gross and microscopic pathology in the calves was consistent with that reported previously (11, 29).

A PCR was used to detect and quantify LSDV genomic DNA in venous blood samples collected from clinical and nonclinical calves. The viraemia of calves from groups A-C have been described previously (18, 50). The viraemia detected in calves from group D is shown in Figures 1E and F. Virus was detected in the blood at 5 dpi in all five inoculated cattle at low levels (approximately 10^2^ genome copies/mL of blood). Virus was detected in the blood of the two nonclinical cattle (D1 and D3) intermittently over the rest of the study period at low levels. In contrast the three clinical calves demonstrated a classic viraemia curve, peaking at between 10^4^ and 10^5^ genome copies/mL at 9 dpi (D2), 13 dpi (D4) or 14 dpi (D5), that decreased by 21 dpi.

These results are consistent with previous studies using this model and showed that a combined intravenous/intradermal inoculation of LSDV results in clinical disease similar to that described in field outbreaks of LSD, characterised by lymphadenopathy, pyrexia, multiple cutaneous lesions and a viraemic curve. Following experimental challenge, only a subset of the challenged calves (41%) developed clinical disease. The nonclinical calves developed a local lymphadenopathy and occasionally a fever spike but did not develop cutaneous lesions distant from the inoculation site and exhibited a low and intermittent viraemia post-inoculation. In order to better characterise the difference between these clinical and nonclinical outcomes we studied the immune response of the intravenous/intradermal inoculated calves.

### LSD is not associated with consistently detectable levels of cytokines in the serum

The immune response of the calves to LSDV inoculation was initially studied by measuring the levels of the pro-inflammatory cytokine IFN-γ and the anti-inflammatory cytokine IL-10 in sera using an ELISA. No IFN-γ or IL-10 were detected in the serum of any of the 17 inoculated calves from studies A-D at any timepoint (data not shown), demonstrating that LSDV challenge does not induce high systemic levels of these cytokines in either clinical or nonclinical presentations.

Type I IFNs (IFN-α and IFN-β) are key anti-viral cytokines (52) therefore the level of type I IFN in the serum of the five calves in group D was measured using a cell-based reporter system. This system uses a modified MDBK cell line (MDBK-t2) containing a MxA promoter driving a chloramphenicol acetyltransferase (CAT) reporter gene (51). Serum samples from 0 to 15 dpi from all five calves in study D were tested. A single peak of type I IFN (75 IU/mL) was detected at 5 dpi in the serum of calf D5, the most severely affected calf in the clinical group (Figure 2A). Type I IFN was not detected in serum samples from nonclinical calves (Figure 2B).

**Figure 2.**
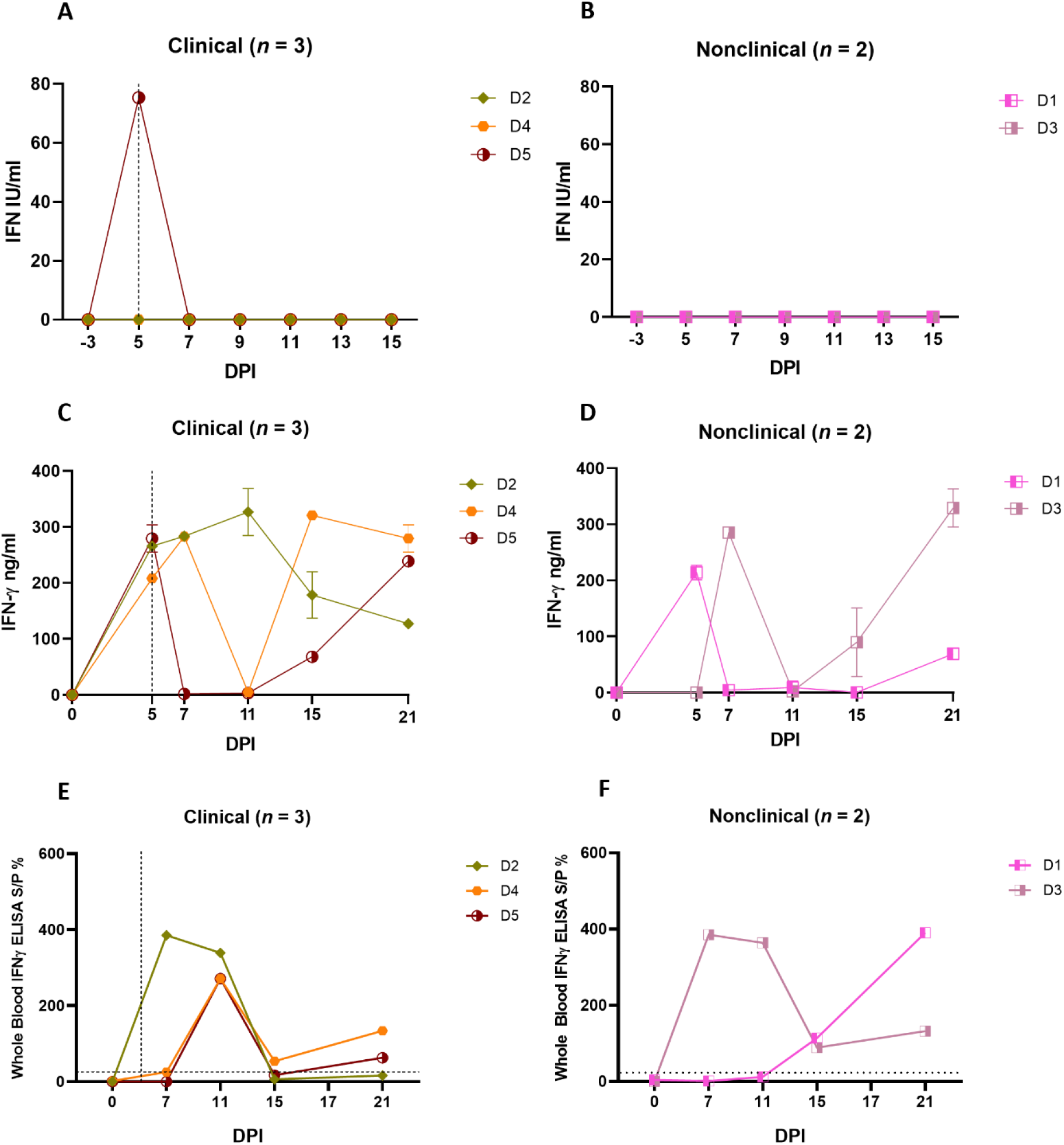
Needle-inoculated clinical and nonclinical calves produce IFN-γ but not type I IFN in response to stimulation of PBMCs or blood by UV-inactivated LSDV. (**A** and **B**) The amount of type I IFN in the sera of needle-inoculated cattle at indicated timepoints post infection was quantified using a cellular reporter system (MxCAT assay). IGRAs were performed on PBMC purified from heparinised blood collected at indicated timepoints from clinical (**C**) and nonclinical (**D**) calves. PBMCs were stimulated overnight with UV-inactivated LSDV, supernatant collected, and IFN-γ quantified using an in-house ELISA. The error bars represent the SEM. An IGRA was also performed on stimulated whole blood from clinical (**E**) and nonclinical (**F**) calves at the timepoints indicated. Whole blood was stimulated with live LSDV overnight, then IFN-γ present in the plasma quantified using a commercially available ELISA. The error bars represent the SEM. The dotted horizontal line represents a 15 S/P % positive cut-off. The dotted vertical line represents the first day cutaneous lesions occurred in the clinical animals. Data corrected to mock PBS stimulation.

### Calves inoculated intravenously and intradermally with LSDV develop a cell-mediated immune response characterised by IFN-γ production by PBMCs following *in vitro* restimulation

In order to measure the CMI response of calves to LSDV, PBMCs from the five group D calves were isolated from whole blood collected at 0, 5, 7, 11, 15 and 21 dpi, and stimulated with UV-inactivated LSDV overnight or PBS as a mock stimulant. The supernatant was then collected and analysed using an in-house ELISA to detect IFN-γ (Figure 2C). No IFN-γ was secreted by the PBMCs in response to LSDV stimulation prior to inoculation at 0 dpi however moderate (100-250 ng/mL) to high (over 250 ng/mL) amounts of IFN-γ were secreted by the PBMCs from calves D2-D5 post-inoculation, indicating a CMI response to intravenous/intradermal LSDV inoculation. This response was first detected 5 or 7 dpi and continued to be expressed inconsistently throughout the study period to 21 dpi. Calf D1 had a lower response with around 100ng/mL IFN-γ detected at 5 and 21 dpi. No clear trend was discerned across the time course of the study, and no difference between the three clinical and two non-clinical calves was observed (Figures 2C and D).

In addition to studying IFN-γ release from purified and stimulated PBMCs, the IFN-γ release assay (IGRA) was also performed on heparinised whole blood in order to assess the potential of this simpler method for detecting the CMI response as a diagnostic test for LSD (Figures 2E and F). Blood collected on 0, 7, 11, 15, 17 and 21 dpi was stimulated overnight with live LSDV, the plasma collected, and IFN-γ quantified using a commercially available ELISA (ID Screen® Ruminant IFN-g ELISA, Innovative Diagnostics) following the manufacturer guidelines. Early peaks in IFN-γ at 7 and 11 dpi were detected in the clinical cattle and decreased by 15 dpi. A similar trend was observed in the nonclinical calf D3 whilst D1 peaked at 21 dpi. No difference was observed between the clinical and nonclinical animals in group D. While both the PBMC and whole blood IGRAs demonstrated the presence of a robust and specific CMI response in all five calves after inoculation with LSDV, there were substantial differences between the IGRA results carried out on PBMCs and whole blood. For example, at 11 dpi, only calf D2 had a strong IFN-γ response when tested using the PBMC IGRA, however calves D2, D3, D4 and D5 all had a strong IFN-γ response at 11 dpi when tested using the whole blood IGRA.

The CMI response was explored further using a bovine IFN-γ ELISpot assay which measures the number of cells producing IFN-γ in response to stimulation. Freshly harvested bovine PBMCs collected from the group D calves at timepoints between 0 and 21 dpi were stimulated overnight with UV-inactivated LSDV, No IFN-γ-producing cells were detected from blood samples collected prior to inoculation. A small number of cells (≤ 100) producing IFN-γ in response to LSDV stimulation were detected in all calves, with the exception of calf D1, as early as 5 dpi, increasing slightly by 7 and 11 dpi (Figures 3A and B). However, the strongest response was seen at 15 and 17 dpi, when up to 2000 spot forming cell (SFC)/10^6^ were detected. This pattern of an increasing frequency of IFNγ producing cells over time was consistent in four of the five calves (D1 and D3-D5) however calf D2 exhibited a different response, with peak IFN-γ producing cells seen at 11 dpi. Similar to the IGRA findings above, there was no obvious difference seen in the results of ELISpot assay between clinical and nonclinical calves in group D.

**Figure 3.**
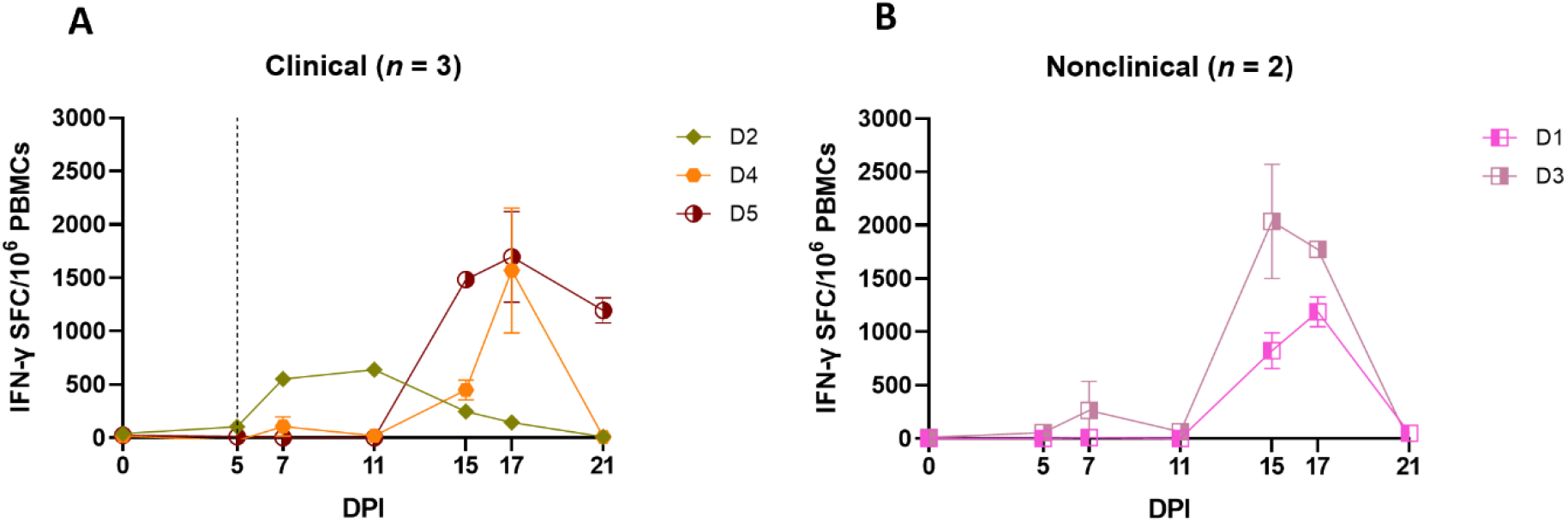
Needle-inoculated clinical (A) and nonclinical (B) calves produce IFN-γ in response to stimulation of PBMCs by UV-inactivated LSDV as measured by ELISpot assay. The number of IFN-γ producing PBMCs stimulated with SW-UV inactivated LSDV was determined by IFN-γ ELISpot and presented as spot forming cells/million (SFC/10^6^). The error bars represent the SEM. The dotted line represents the first day of the onset of cutaneous lesions in the clinical animals. Data corrected to mock PBS stimulation.

In order to determine which subpopulation of PBMCs was responsible for the IFN-γ release seen in the IGRA and ELISpot assays, an intracellular cytokine staining (ICS) assay was performed on PBMCs freshly isolated from the five inoculated calves on 0, 7, 11, 15 and 21 dpi. The PBMCs were stimulated overnight with UV-inactivated LSDV followed by surface labelling of cell surface markers CD4 and CD8β and intracellular labelling of IFN-γ. Flow cytometry was used to determine the percentage of CD4^+^CD8^-^, CD4^-^CD8^+^ or CD4^-^CD8^-^ cell populations producing IFN-γ in response to LSDV stimulation (Figure 4A - F). At 0 dpi, IFN-γ was not detected in any of the cell types investigated. In all the cattle, irrespective of their clinical status, low levels (< 0.6 %) of CD4^-^CD8^-^ cells produced IFN-γ at any of the subsequent timepoints investigated (Figure 4E-F). The CD4^-^CD8^+^ populations of the two non-clinical calves (D1 and D3) and one clinical calf (D2) generated weak IFN-γ responses, not exceeding 0.5 % of the population, throughout the study. In contrast, IFN-γ responses were detected in the CD4^-^CD8^+^ populations of two of the clinical calves (D4 and D5) at 15 dpi, when 1.7 % and 0.5 % of the CD4^-^CD8^+^ populations produced IFN-γ in response to LSDV stimulation. In the CD4^+^CD8^-^ population, low to moderate level responses were observed from 7 dpi onwards in D1, D2, and D3, with the strongest response in the nonclinical calves D1 and D3 detected at 21 dpi. Very low responses were detected in the CD4^+^CD8^-^ population in the clinical calves (D4 and D5) prior to 15 dpi, with the strongest IFN-γ response of 3.65% and 3.8 % of the CD4^+^CD8^-^ population detected at 21 dpi in calves D4 and D5 respectively. Overall, production of IFN-γ by both CD4^+^CD8^-^ and CD4^-^CD8^+^ T cells in response to stimulation with LSDV was detected in calves D1, D3, D4 and D5, with the CD4^+^CD8^-^ cell population having a stronger response at later time points. Inter-calf variation was present, with calf D2 producing a more muted response in both CD4^+^CD8^-^ and CD4^-^CD8^+^ cell populations. Very weak responses were observed in the CD4^-^CD8^-^ cell population across all five cattle at all five time points.

**Figure 4.**
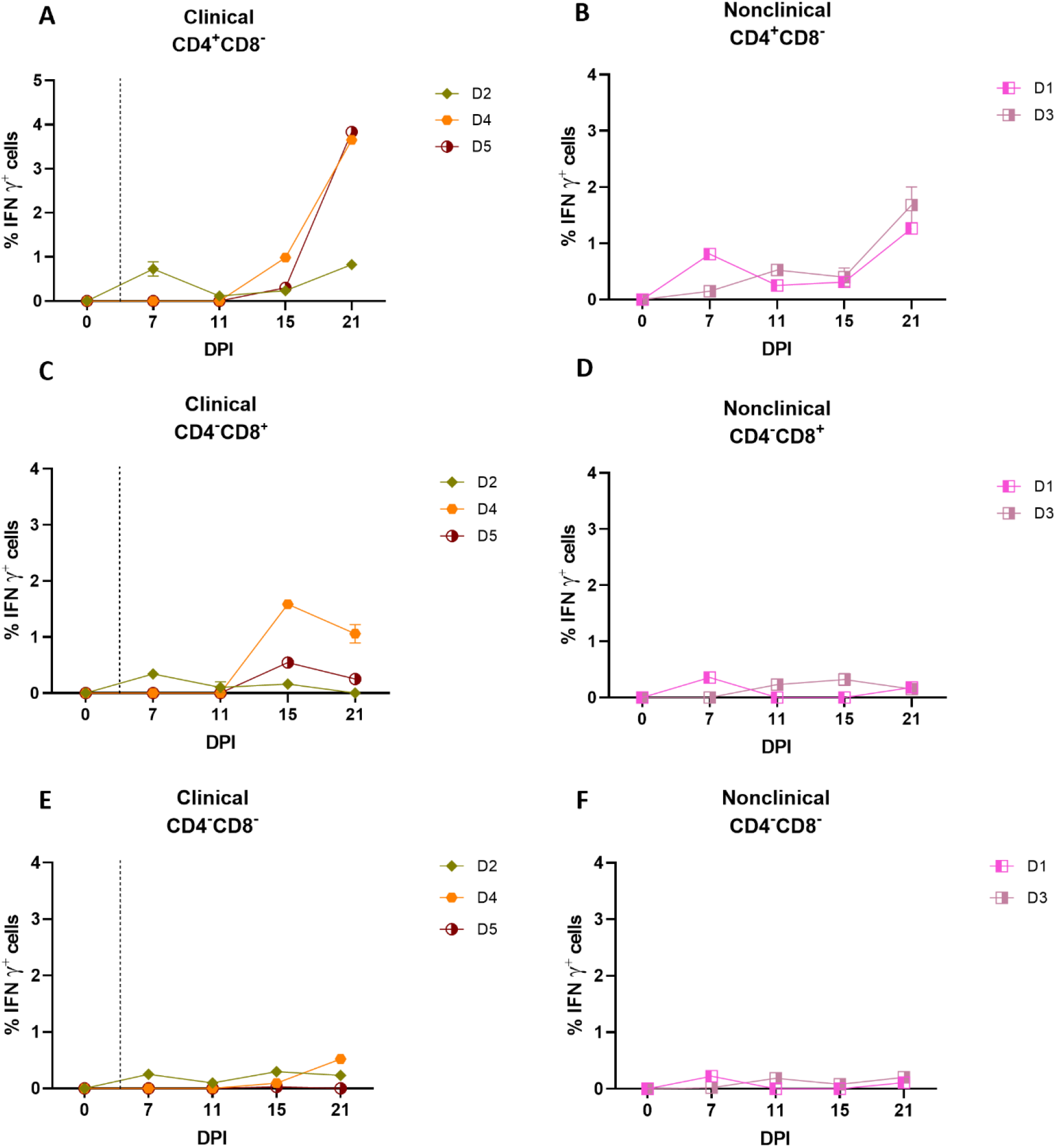
CD4^+^CD8^-^ and CD8^+^CD4^-^ but not CD4^-^CD8^-^ T cells are responsible for the production of IFN-γ in response to stimulation with UV-inactivated LSDV in needle-inoculated calves. Flow cytometric analysis was used to determine the % of IFN-γ^+^ T cells expressing CD4^+^CD8^-^ (**A** and **B**), CD8^+^CD4^-^ (**C** and **D**) and CD4^-^CD8^-^ (**E** and **F**) in response to stimulation with SW-UV inactivated LSDV. Samples were tested in triplicate and normalised to the mock control. The error bars represent the SEM. The dotted vertical line represents the first day lesions occurred in clinical animals. Data corrected to mock PBS stimulation.

In summary, the IGRA, ELISpot and ICS assays indicate that calves inoculated with LSDV intravenously and intradermally develop an IFN-γ response by 5-7 dpi. Both clinical and nonclinical calves developed a CMI with no distinguishing features particular to either group identified.

### Clinical and nonclinical calves inoculated with LSDV develop a detectable humoral immune response

The antibody response to intravenous/intradermal LSDV challenge in calves from groups A-D was initially studied using a commercial ELISA. Six of the 7 clinical calves from groups A to D (A3, A5, B9, C12, D2, and D4) were positive by ELISA for bAbs by 21 dpi (Figure 5A). Animals A5, B9 and C12 were the first to reach the positive cut-off point at 15 dpi. The seventh clinical animal, calf D5, remained negative but a rising antibody level below the cut-off S/P % was detected at 17 and 21 dpi. There were 10 nonclinical calves in Groups A to D (including 3 negative control animals A1, B6 and C11). Calf B7 was the only to develop a positive ELISA result, on days 15 and 17 dpi (Figure 5B) with calves D1 and D3 borderline positive by 21 dpi. These results suggest that clinical calves develop a more rapid and robust humoral immune response compared to nonclinical calves following intravenous/intradermal inoculation.

**Figure 5.**
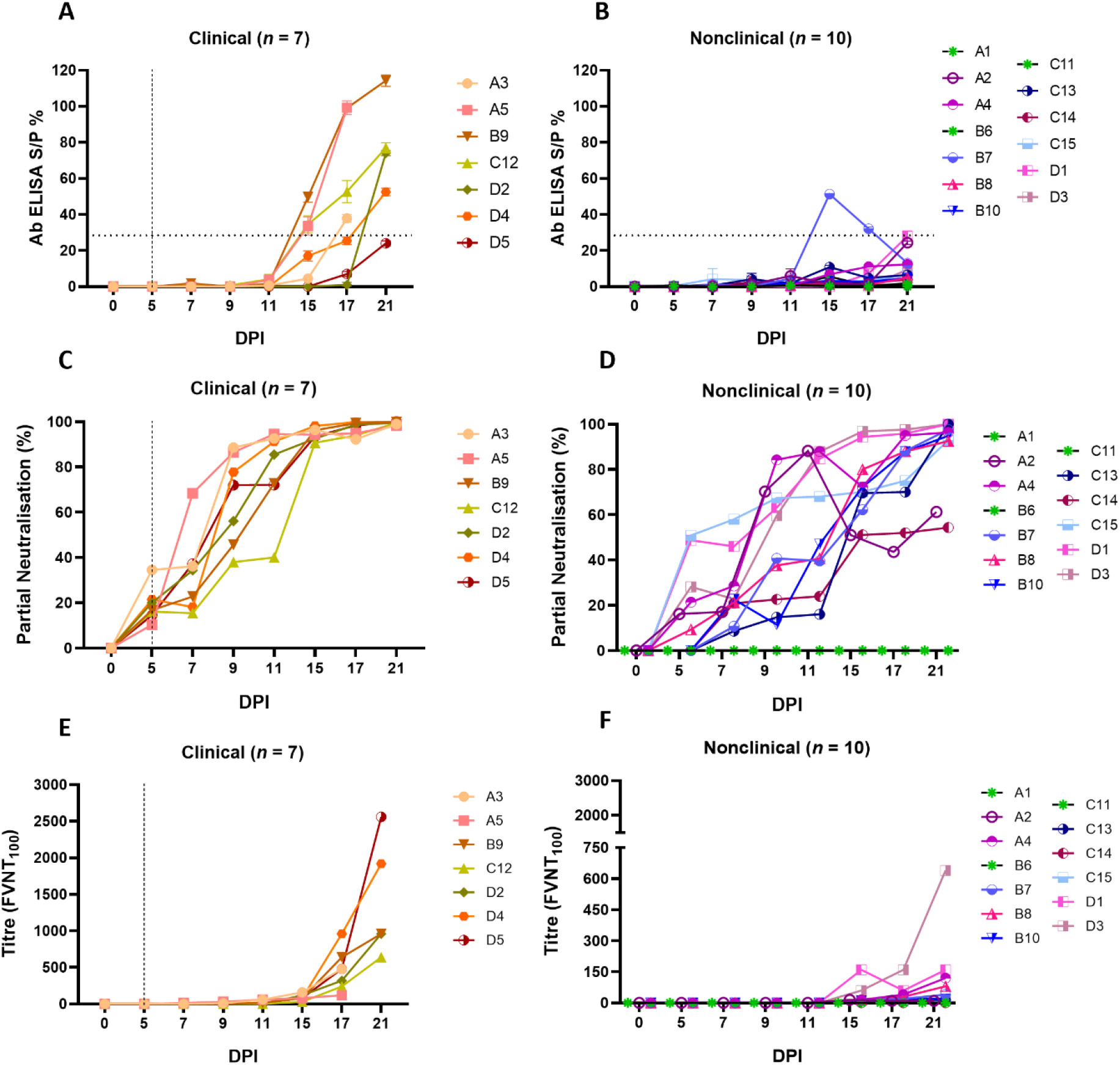
Calves that develop clinical disease following needle-inoculation with LSDV also develop a more rapid and robust humoral immune response as measured either by ELISA or FVNT. **The a**ntibody responses in calves after needle-inoculation with LSDV was measured using the ID Screen® Capripox Double Antigen Multi-species ELISA kit (Innovative Diagnostics) (**A** and **B**). The horizontal line represents the 30 S/P% cut-off. The production of neutralising antibodies was measured using a fluorescent virus neutralisation test. The percentage of viral foci-forming units neutralised by the sera was measured over time in clinical (**C**) and nonclinical (**D**) calves. Complete neutralisation of the virus (FVNT_100_) was also calculated in clinical (**E**) and nonclinical (**F**) calves. The dotted line represents the first day of the onset of cutaneous lesions in the clinical animals.

In order to quantify functional antibodies, a FVNT was performed to detect neutralising antibodies (nAbs) to LSDV. This assay takes advantage of the “foci” formed by LSDV on MDBK cells (49). Monitoring the reduction in the number of foci at each dilution was used as an indicator of neutralising activity (using a cut-off of 50 foci/well), enabling partial neutralisation to be quantified over the duration of each study. Evidence of neutralising activity was detected in the serum of all 7 clinical calves at 5 dpi, with values between 10.37 – 34.50 % (Supplementary Table 1). The degree of neutralisation then increased over time until all clinical animals had antibodies that were able to neutralise 100% of LSDV at 17 and 21 dpi (Figure 5C). There was moderate variation between individual animals with calf C12, for example, exhibiting a slower rate of increase over time. In contrast, pronounced variation existed in the partial neutralisation activity of serum from the calves in the nonclinical group (Figure 5D). At 5 dpi, antibodies from 4 of the 10 nonclinical calves did not exhibit any detectable neutralisation (B7, B10, C12 and C14), while antibodies present in the remaining calves neutralised between 9.25 and 50.88% (Supplementary Table 2). of LSDV. Different rates of increase in partial neutralisation over time were evident between all nonclinical calves. At 11 dpi antibodies from calves A2, A4, C15, D1 and D3 neutralised 68 – 88% of LSDV and with antibodies present in the remaining calves neutralised 16 – 47%. At 21 dpi calves, A2 and C14 neutralised 54 – 61% of LSDV whilst antibodies in the remaining animals neutralised 95 – 100%.

Complete neutralisation determined by FVNT_100_ based on the gold standard test for nAbs, as described in the WOAH Terrestrial Guide for capripoxviruses (53), was observed in all clinical and nonclinical animals except for nonclinical calf C14. The clinical calves generated higher titres of nAbs by 21 dpi (120 – 2560; Figure 5E, Supplementary Table 2) compared to the nonclinical calves (10 – 160 except for calf D3 with a titre of 640; Figure 5F; Supplementary Table 2). Moderate variation in nAb titres between individual clinical calves was evident.

In conclusion, detailed analysis of the immune response of calves inoculated via the intravenous/intradermal route identified a rapid (5-7 dpi) cell-mediated and humoral immune response to LSDV. There were no consistent differences in the cell-mediated immune response between the clinical and nonclinical calves, however a stronger and more rapid humoral immune response was detected in clinical animals compared to the nonclinical animals.

### Arthropod inoculation

#### Inoculation of calves with virus-positive insects results in development of clinical disease

The predominant means of LSDV transmission in the field is via haematogenous arthropods including flies and mosquitoes (17, 19, 20). In order to determine whether the method of inoculation influences the immune response of calves to LSDV, *S. calcitrans* flies and *Ae. Aegypti* mosquitoes were partially fed on cutaneous nodules from a clinical calf (calf D5), and allowed to immediately (within 1 h) re-feed on naive calves in group RS and RA respectively (Table 1). The feeding (from donor group D calves) and refeeding (on recipient RS and RA calves) occurred on five consecutive days (0-4 dpi), with 20 *S. calcitrans* flies or *Ae. Aegypti* mosquitoes fed on each calf in the RS and RA groups each day, to give a total of 100 arthropods feeding on each RS and RA calf. Arthropods were fed on the paravertebral dorsum of the RS and RA calves, on either side of the vertebral column between the scapula and tail. Animals in groups RS and RA were monitored for clinical signs of LSD throughout the study period.

Four of the five RS calves (RS1, RS2, RS4 and RS5) and four of the five RA calves (RA1, RA3, RA4 and RA5) developed clinical LSD (cutaneous lesions) following arthropod inoculation. Severe disease (>100 cutaneous lesions per animal) was evident in calves RA1, RA3, RA5, RS1, RS2 and RS4 (Figure 6A), moderate disease in calf RA4 (50 lesions) and mild disease was observed in RS5 (1 lesion). Lesions were first noted on 11 (RA1, RA3, RA4 and RA5), 12 (RS1, RS2 and RS4) or 13 (RS5) dpi. No cutaneous lesions were detected in the nonclinical calves (Figure 6B). An increase in body temperature was detected in all eight clinical calves from 11 dpi (Figure 6C), persisting for between 1-12 days. A single day of raised temperature was observed in nonclinical calf RA2 at 13 dpi. No temperature increase was detected in nonclinical calf RS3 (Figure 6D). Reddened and swollen inoculation sites, progressing to necrosis of the skin, were noted in the clinical calves with the exception of RS5. Increased size of the prefemoral and prescapular lymph nodes was present in all ten calves from 10-14 dpi onwards, lasting in some calves to the end of the study. This lymphadenopathy was more marked in the clinical calves. Loss of body condition score was noted in all clinical calves except RS5. Lethargy was noted in calf RA1 from 15-16 dpi, RA3 on 18 dpi, and RS4 15-20 dpi. Calves RA1, RA3 and RS4 were euthanised at 21 dpi, RS1, RS2, RA4 and RA5 at 25 dpi and calves RS3, RS5 and RA2 at 29 dpi.

**Figure 6.**
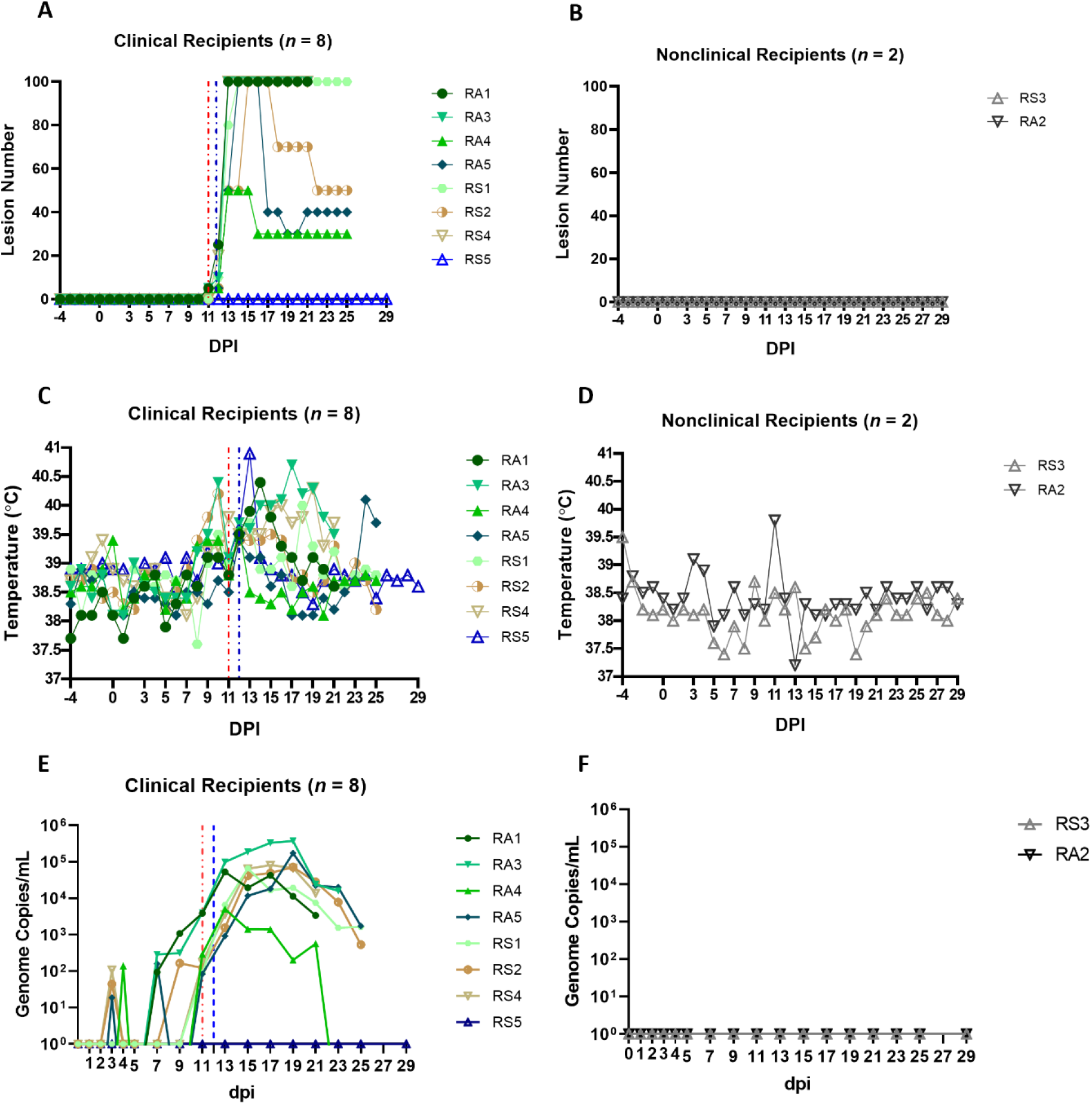
Eight out of ten arthropod-inoculated calves developed clinical signs consistent with LSD approximately 11 days after the start of inoculation. The number of cutaneous lesions in the clinical (**A**) and nonclinical (**B**) calves were recorded each day, up to a maximum of 100 lesions. The rectal temperature of clinical and nonclinical calves (**C** and **D**) was recorded daily. The number of LSDV genome copies in the blood of each calf was quantified by qPCR. The red dotted line represents the onset of cutaneous lesions for group RS cattle and the blue dotted line the onset of cutaneous lesions for group RA cattle.

LSDV viraemia (genome copies/mL detected by qPCR) was detected as early as 3 dpi in arthropod inoculated calves (Figure 6E). Seven clinical calves with moderate or severe disease (RA1, RA3, RA4, RA5, RS1, RS2, and RS4) exhibited a similar viraemia curve with peak viraemia occurring at 17-19 dpi. Calves RA3 and RA5 had the highest viraemia levels at 3.7 × 10^5^ and 1.7 × 10^5^ genome copies/mL. Calf RA4 which developed moderate disease had a shorter and lower magnitude viraemia when compared to the severely affected calves. Calf RS5 developed mild clinical disease with only a single cutaneous lesion. The nodule contained high levels of LSDV genomic DNA (1.4 × 10^5^ genome copies of LSDV/mL of skin microbiopsy extract), confirming the pathology was caused by LSDV. This calf remained viraemia negative throughout the study across all 17 time points that were blood sampled. It was clinically inapparent on other parameters scored throughout the study apart from 13 dpi when it had a rise of 1.3°C in temperature to 40.9°C, and a high heart rate of 90 beats per minute. These parameters had returned to normal by 14 dpi. The nonclinical calves RA2 and RS3 were negative for LSDV viraemia throughout the study.

In summary, the clinical signs, gross pathology noted in the arthropod-inoculated calves was consistent with that reported in “needle inoculated” calves including those challenged via the intravenous/intradermal route. The magnitude and length of viraemia was also similar, however the preclinical or latent period was longer in the arthropod-inoculated calves (11-13 dpi in groups RS and RA compared to 5-6 dpi in groups A-D).

### Type I IFN can be detected intermittently in the serum of calves inoculated with virus-positive insects

We measured IFN-γ and IL-10 in the sera of the RS and RA calves but, similar to the calves in group D, no cytokine expression was detected at any of the timepoints tested (data not shown). We then used the MxCAT assay to look for type I IFN in the sera and detected this cytokine in both RS and RA groups across the time course of the study (Figure 7A). Seven clinical recipient animals exhibited a type I IFN response. Five of these calves were characterised as severely diseased (RA1, RA3, RA5, RS1, RS2), one as moderate (RA4), and one (RS5) exhibited only mild signs of LSDV infection. Most of these calves demonstrated a single peak of type I IFN between 9 and 15dpi, ranging between 4.3-191 IU/mL, except calf RS5 which had 191 IU/mL of type I IFN at 13dpi and 16.5 IU/mL at 15dpi. No type I IFN was detected in the serum of the two nonclinical calves RA2 and RS3 or the severely affected calf RS4 at any time point tested.

**Figure 7.**
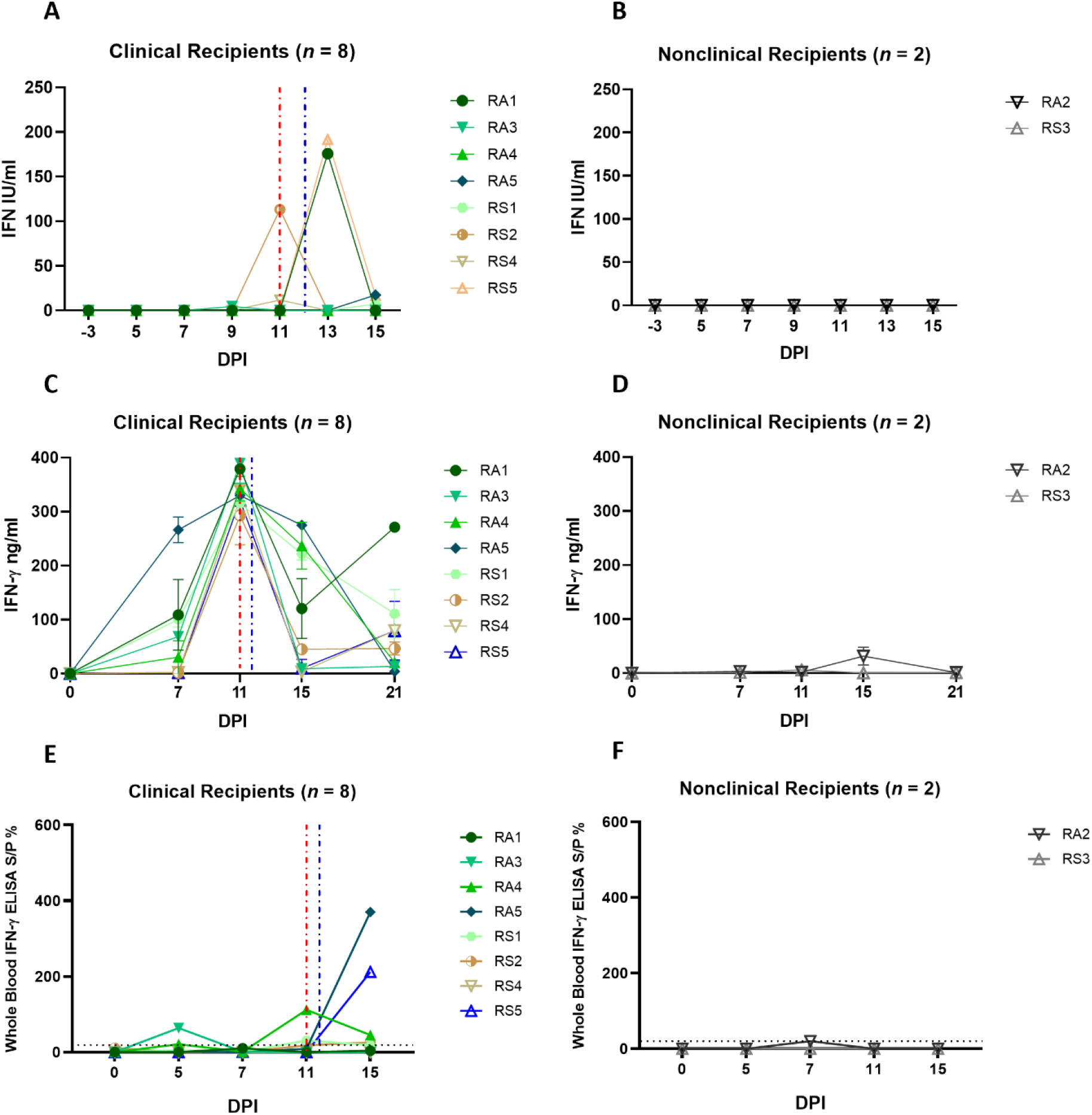
Arthropod-inoculated clinical calves but not nonclinical calves develop a uniform and robust LSDV-specific CMI response as measured by IGRA on purified PBMCs. The amount of type I IFN in the sera of needle-inoculated cattle at indicated timepoints post infection (**A** and **B**) was quantified using a cellular reporter system (MxCAT assay). IGRAs were performed on PBMC purified from heparinised blood collected at indicated timepoints from clinical (**C**) and nonclinical (**D**) calves. PBMCs were stimulated overnight with UV-inactivated LSDV, supernatant collected, and IFN-γ quantified using an in-house ELISA. The error bars represent the SEM. An IGRA was also performed on stimulated whole blood from clinical (**E**) and nonclinical (**F**) calves at the timepoints indicated. Whole blood was stimulated with live LSDV overnight, then IFN-γ present in the plasma quantified using a commercially available ELISA. The error bars represent the SEM. The dotted horizontal line represents a 15 S/P % positive cut-off. Red dotted line represents the onset of lesions for group RS cattle and the blue dotted line for group RA cattle. Data corrected to mock PBS stimulation.

### A CMI response to LSDV can be detected in clinical but not nonclinical calves after inoculation with virus-positive arthropods

PBMCs isolated from blood collected from recipient calves on 0, 7, 11, 15 and 21 dpi were stimulated with UV-inactivated LSDV and an IGRA performed as described above. A strong peak in IFN-γ secretion (250 – 380 ng/mL) was detected at 11 dpi in all eight clinical recipient calves (RS1, RS2, RS4, RS5, RA1, RA3, RA4, RA5), with lower levels at 15 and 21 dpi (Figures 7C and D). In contrast very low levels of IFN-γ (<50 ng/mL) were produced in response to LSDV stimulation in the two nonclinical calves RS3 and RA2 at all time points. Both the RA and RS group show a consistent trend amongst the clinical cattle with a peak of CMI at 11 dpi, in contrast to the very low levels of IFN-γ produced by the two nonclinical calves.

The IGRA assay was also carried out on whole blood, using live LSDV as overnight stimulant. Lower levels of IFN-γ secretion were detected using this method in all clinical animals compared to the response seen in stimulated PBMCs with convincing responses detected only in calf RA3 at 5 and 11 dpi and RA5 and RS5 at 15 dpi (Figure 7E). Low or no IFN-γ was detected in the two nonclinical calves RS3 and RA2 at any timepoint. The lack of consistency between the PBMC and whole blood IGRA results reflects that seen in the analysis of the group D responses.

Overall, these data indicate that there is a strong anti-LSDV CMI response present in all eight arthropod inoculated calves at 11 dpi, at or prior to the time they develop cutaneous lesions. The absence of a detectable CMI response in the two nonclinical calves suggest early local antiviral defences were sufficient to control the virus in these animals, avoiding the need for a systemic response.

### Clinical but not nonclinical calves develop strongly neutralising antibody titres after inoculation with virus-positive arthropods

Serum samples were evaluated for bAbs specific to LSDV by commercial ELISA. Clinical calves RS4 and RA1 showed the presence of bAbs at 21 dpi, and calves RS2 and RA5 were positive at 25 dpi (Figure 8A). The remaining clinical calves RS1, RS5, RA3 and RA4 were negative as were the three nonclinical calves (RS3 and RA2; Figure 8B), although rising titres of bAbs were visible at later time points.

**Figure 8.**
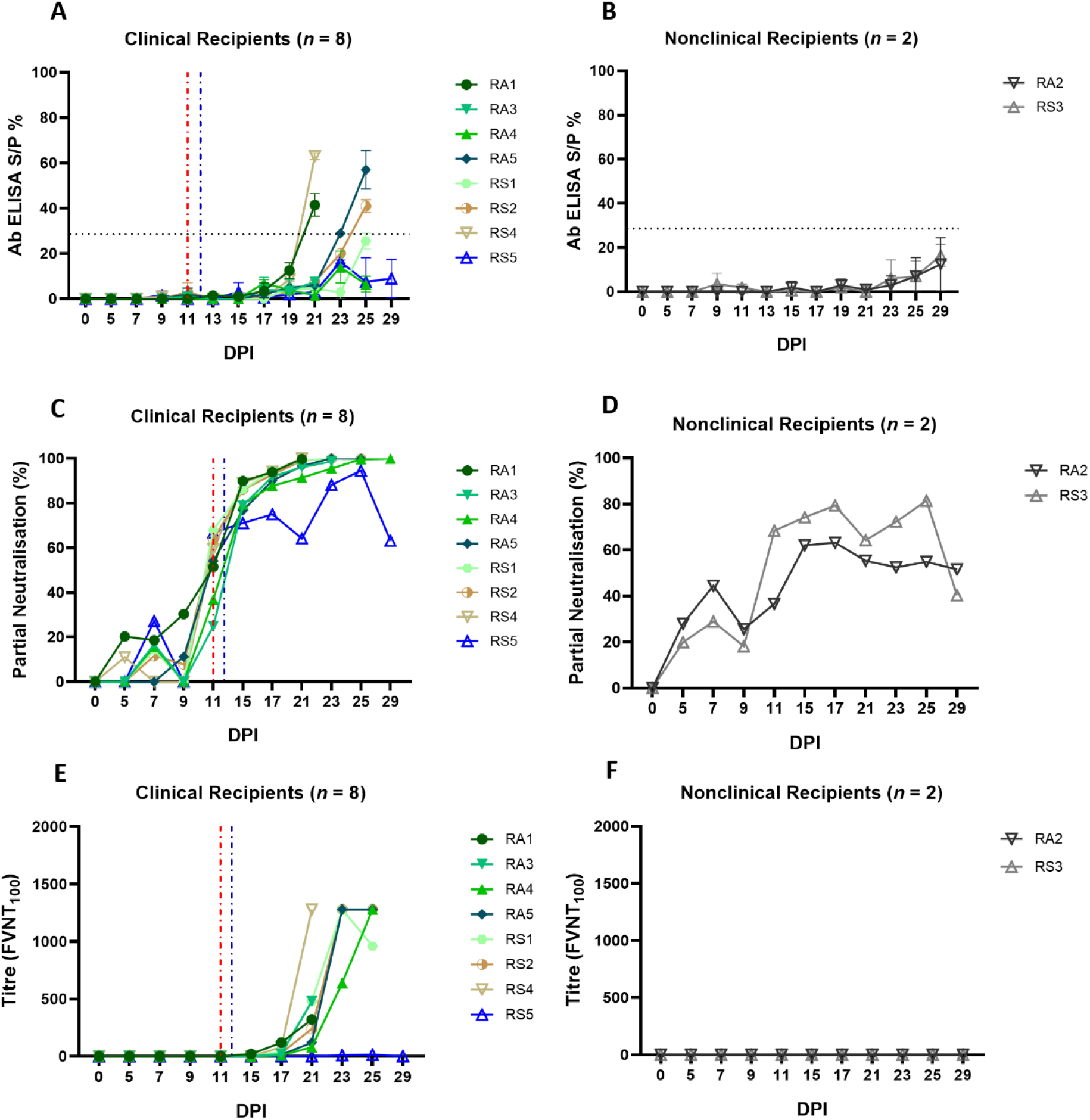
Calves that develop clinical disease following arthropod inoculation with LSDV develop a rapid and robust humoral immune response as measured either by ELISA or FVNT. The antibody responses in calves after needle-inoculation with LSDV was measured using the ID Screen® Capripox Double Antigen Multi-species ELISA kit (Innovative Diagnostics) following manufacturer instructions (**A** and **B**). The error bars represent the SEM. The production of neutralising antibodies was measured using a fluorescent virus neutralisation test. The percentage of viral foci-forming units neutralised by the sera was measured over time in clinical (**C**) and nonclinical (**D**) calves. Complete neutralisation of the virus (FVNT_100_) was also calculated in clinical (**E**) and nonclinical (**F**) calves. Red dotted line represents the onset of lesions for group RS cattle and the blue dotted line for group RA cattle.

The FVNT was used to detect and quantify neutralising antibodies against LSDV in the sera of the arthropod challenged calves. No nAbs were detected in the sera of any of the calves at 0 dpi, however, partially nAbs were generated by all ten calves during the study. Seven of the eight clinical calves exhibited a rapid increase in partial nAbs from 11 dpi onwards (24.87 – 68%; Supplementary Table 3) until 100 % neutralisation was observed from 15 dpi (Figures 8C). Mildly affected calf RS5 had a slower increase in nAbs first detected at 7 dpi, to peak just below 100% at 25 dpi before declining at 29 dpi. Nonclinical calves RS3 and RA2 had low-level partial nAbs detected at 7 dpi (27.25 – 44.50%) that increased to peak first at 17 dpi or 25 dpi before declining by 29 dpi (Figure 8D; Supplementary Table 3).

Complete neutralisation titres were detected in all clinical calves (with the exception of RS5) starting from 15 dpi for calf RA2 with an FVNT_100_ titre of 20 (Figure 8E; Supplementary Table 4). Final titres generated at individual animal endpoints were between 320 – 1280. No complete neutralisation titres were detected in the two nonclinical calves (Figures 8F; Supplementary Table 4).

In summary, all 10 arthropod inoculated calves developed a detectable antibody response to LSDV. This indicates that the arthropod challenge was successful at exposing all ten calves to LSDV. The magnitude of the humoral response correlated well with the severity of disease, with the severe and moderately affected calves rapidly developing neutralising antibodies capable of completely neutralising the input virus. The mildly affected calf RS5 and the two nonclinical calves developed lower titres of nAbs.

### The B cell response of nonclinical calves post-challenge with virus-positive insects is characterised by a strong IgM response and delayed class switching

In order to investigate the humoral immune response to LSDV in more detail, the expression of LSDV-specific IgM and IgG antibodies was examined using a B cell ELISpot. This was performed on PBMCs to quantify the number of LSDV-specific antibody secreting cells (ASCs). In group D (needle inoculated calves) an initial peak in IgM ASCs (247.4 ± 36.96 ASCs/10^6^ PBMCs) was detected in both clinical and nonclinical calves at 5 dpi, around the onset of the development of skin lesions. The number of IgM ASCs decreased slightly at 7 dpi and peaked again at 15 dpi (466.30 ± 53.56 ASCs/10^6^ PBMCs; Figures 9A and B). The predominant isotype detected at 21 dpi was IgG suggesting a class switch to IgG. In the clinical calves an average of 224.3 ± 130.9 ASCs/10^6^ PBMCs was determined. A similar pattern of LSDV-specific ASC quantification was seen in the two nonclinical calves, with an increase in IgG and a decrease in IgM at 21 dpi. There was no clear difference in the clinical and nonclinical calves in group D with respect to the ratios of IgG and IgM and potential class switching.

**Figure 9.**
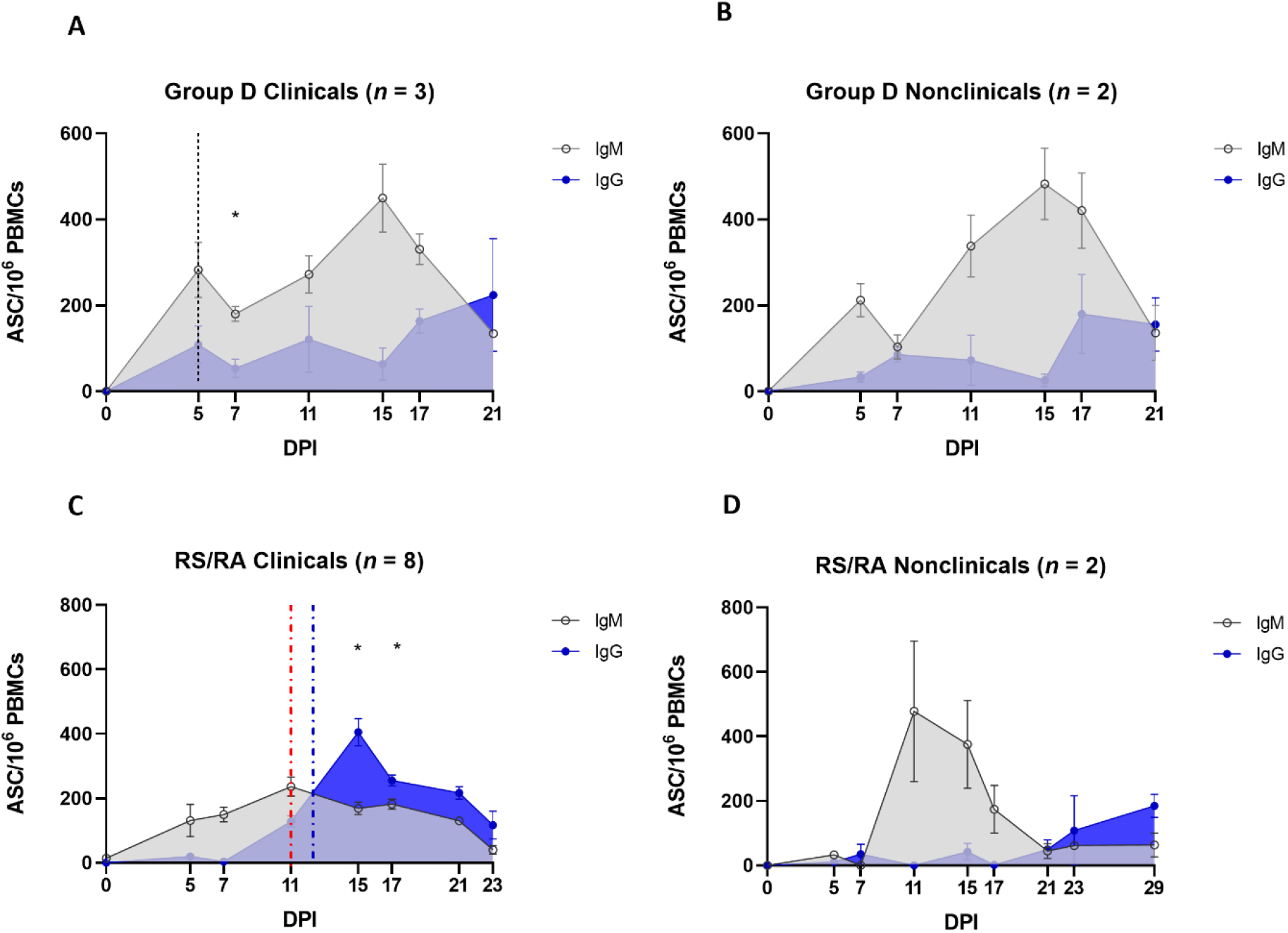
Protection against LSD is associated with a robust LSDV-specific IgM response following arthropod-inoculation but not needle-inoculation. PBMCs were isolated from heparinised blood from needle-inoculated (**A** and **B**) and arthropod-inoculated (**C** and **D**) calves at the time points shown, and stimulated overnight with live LSDV. Antibody secreting cells were then labelled with bovine anti-IgM or bovine anti-IgG antibodies. Spots were read using an ImmunoSpot 7.0 reader and ImmunoSpot SC suite (Cellular Technology Limited). Results were manually validated for false-positive results and expressed as the mean number of ASCs/million. Results are expressed as group mean ASCs/10^6^ PBMCs ± standard error of the mean. Red dotted line represents the onset of lesions for group RS cattle and the blue dotted line for group RA cattle. Data corrected to mock uninfected cell lysate stimulation.

In comparison, the pattern of detection of IgM and IgG secreting ASCs in arthropod inoculated calves was very different. When compared to the calves in group D, a much lower number of LSDV-specific IgM producing ASCs were present in the PBMCs from clinical recipient calves over the entire sampling period (compare Figure 9A to 9C). A slow increase over the first 11 dpi reached a plateau between 11 and 17 dpi (151.8 ± 9.14 ASCs/10^6^ PBMCs) and then reduced until 23 dpi when no LSDV-specific IgM ASCs were detected in calves RS1, RS2, RA4 and RA5 (Figure 8C). There was a rapid increase in LSDV-specific IgG ASCs from 7-15 dpi to a peak of 405.4 ± 42.4 ASCs/10^6^ PBMCs at 15 dpi in the clinical calves. This resulted in a significant change from a predominance of IgM to IgG at 15 dpi (*P* = 0.0356) and 17 dpi (*P* = 0.0367). The number of IgG ASCs specific for LSDV then decreased over the subsequent sampling time points until the end of the study. The IgM and IgG secreting ASC profiles from the nonclinical recipient calves were very different to the clinical calves. The PBMCs from these two calves (RA2 and RS3) contained numerous LSDV-ASCs secreting LSDV-specific IgM by 11 dpi (575 ± 115.20 ASCs/10^6^ PBMCs). The number of ASCs secreting LSDV-specific IgM then declined by 21 dpi (Figure 8D), with a concurrent slow increase in LSDV-specific IgG-secreting ASCs leading to a predominance of IgG ASCs by 23 dpi.

Overall, these data have shown the rapid (by 5 dpi) appearance of IgM-secreting LSDV-specific ASCs in calves inoculated via the intravenous/intradermal route, followed by a slower increase in IgG-secreting LSDV-specific ASCs. This profile of ASC is not influenced by the clinical outcome of inoculation. However, the timing and magnitude of LSDV-specific ASC production in arthropod challenged calves is dependent on clinical outcome. Nonclinical calves contain more LSDV-specific IgM ASCs than clinical calves, especially at 11 and 15 dpi when multifocal cutaneous nodules were first appearing on calves in the clinical cohort. Fewer IgM-secreting and more IgG-secreting LSDV-specific ASCs in the clinical calves led to an earlier change in the ratio of IgM to IgG suggesting class switching occurs more rapidly in clinical calves (by 15 dpi) when compared to the nonclinical calves (23 dpi).

## 4 Discussion

This study characterises and integrates the clinical, virological, and immunological response of calves following inoculation with LSDV using a needle-inoculation method or via virus-positive arthropods. The use of arthropods to inoculate calves with LSDV was particularly important as this is a clinically relevant route of transmission. Both routes of inoculation produced clinical disease similar to that observed in the field, characterised by pyrexia, lymphadenopathy, and multiple cutaneous nodules, however there were differences noted between the two inoculation methods including the length of the incubation period preceding disease onset, the kinetics of the cell-mediated and humoral immune responses, and the ability to discriminate between clinical and nonclinical calves using immune signatures.

The two inoculation routes deliver very different doses of virus to different compartments of the immune system which may explain the differences. The ID/IV needle inoculation delivered 1 × 10^6^ PFU of LSDV into the skin and 2 × 10^6^ PFU directly into the vein where it would have direct access to the splenic immune compartments, bypassing lymph nodes. In comparison the maximum estimated dose delivered by the arthropods to calves in the RS and RA groups was 3 × 10^3^ PFU per calf. This dose would have been delivered into the dermis, activating local dermal immune responses before draining to regional lymph nodes. The arthropod delivery was also extended over 5 days in order to mimic a field situation more accurately, rather than being delivered via needle inoculation in a single dose. The blood-feeding arthropods would also have delivered virus accompanied by biting trauma and saliva inoculation. The feeding behaviour of arthropods has been shown previously to influence the outcome of virus inoculation (17, 54, 55), although no work has been published on the impact of arthropod feeding on LSD.

Research into other poxvirus models has shown that dose can impact the latent period. Mice inoculated with 10^3^ PFU of vaccinia virus via an intradermal route developed lesions at 6 dpi but later (8 dpi) when the dose was lowered (56). The influence of viral dose was studied in cattle inoculated intravenously with four different doses of LSDV (approximately 3 × 10^2^, 3 × 10^4^, 3 × 10^6^ or 3 × 10^7^ cell culture infectious dose_50_ (CCID_50_)). One animal that was inoculated with the lowest dose displayed a delayed onset of clinical signs (19 dpi) compared to other cattle in the study (7-9 dpi) (57). The incubation period of 11-12 dpi seen in our study in the RS and RA groups is consistent with previous studies which reported incubation periods following arthropod inoculation of LSDV of 12-26 dpi (19), and 10-14 dpi (20). This data suggests that the incubation period of LSD in the field is longer than the 5-7 days reported in experimental studies using needle inoculation, and more likely to be 10-14 days and possibly longer.

The immune response of calves to LSDV inoculation was initially studied by examining the levels of cytokines IFN-γ and IL-10 in the serum using ELISA. Neither of these two cytokines were detected in calves following either needle or arthropod inoculation, suggesting that the levels of systemic cytokines are not substantially raised in LSD, and that detection of a cytokine signature is unlikely to be of diagnostic value in the field. Interestingly, type I IFN was detected in the serum of cattle following LSDV inoculation. In group D only calf D5 had detectable type I IFN, and only at one time point (5 dpi), while type I IFN was detected in the serum of mild and severely affected calves in the RS and RA groups between 9-15 days after arthropod inoculation. No type I IFN was detected in the nonclinical calves. The pattern of type I IFN in LSD was unusual, as a single strong peak of up to 191 IU/mL, with the timing of the peak closely associated with an increase in levels of virus in the blood and the appearance of the cutaneous lesions (5 dpi for calf D5, and between 9-13 dpi for the calves in groups RS and RA). In comparison, the production of type I IFN response to BVDV and FMDV has been shown to be more rapid, of lower magnitude and longer lived. BVDV inoculation resulted in a gradual increase then decrease in levels of type I IFN in the serum from 0-7dpi in animals infected with type 1 and type 2 BVDV (58), with the levels of type I IFN ranging from 5-75 IU/mL. Cattle infected with FMDV showed a peak of serum type I IFN at around 2dpi estimated between 3-6 IU/mL, decreasing to low levels at 7dpi (58). Our study reveals that LSD causes a brief spike in systemic type I IFN closely associated with the onset of skin lesions in clinical calves. This suggests that LSDV is able to suppress the type I IFN response very effectively throughout the course of disease, except at the onset of systemic pathology.

A cell-mediated immune response specific to LSDV was consistently seen in both clinical and nonclinical needle-inoculated calves in group D from the earliest timepoints examined (3 or 5 dpi). This response was studied using IGRAs, ELISpot and ICS which collectively built a picture of a small population of PBMCs producing a high amount of IFN-γ at 5 and 7 dpi, and a larger population of cells generating IFN-γ later in the course of the disease. Both CD4^+^ and CD8^+^ T lymphocytes produced IFN-γ in response to LSDV stimulation, particularly at the later time points (15 and 21 dpi), but the CD4^-^CD8^-^ population of T lymphocytes did not. These results are consistent with previous experimental studies describing the CMI response to LSDV (22, 32, 45). We found that the CMI response of calves following intradermal and intravenous inoculation varied substantially between animals with calf D2 in particular showing a different timing and magnitude of response across IGRA, ELISpot and ICS when compared to the other four calves in the group. It may be that the large intravenous LSDV bolus generated an exaggerated, non-specific, and poorly controlled CMI response in the needle-inoculated calves. Crucially, even though we used multiple methods to characterise the CMI response and examined timepoints through the course of disease, we were unable to differentiate between the CMI response of clinical and non-clinical calves.

In contrast, examination of the CMI response in arthropod-inoculated calves using the IGRA revealed a clear and consistent difference between the clinical and non-clinical cattle. All 8 clinical calves generated a strong and remarkably uniform CMI responses at 11 dpi, with increasing and decreasing amounts of IFN-γ detected at 7 and 15 dpi, respectively. No CMI response was detected in the two nonclinical calves. This may be because the nonclinical calves were able to control the virus at a local level and therefore did not need to activate a systemic CMI response. The very consistent CMI response seen in the arthropod-inoculated calves was surprisingly given the variable dose most likely received by each animal from the virus-positive arthropods, and the 5-day period over which the dose was given. Despite these variables, the calves developed a very uniform response. Interestingly, a number of the clinical arthropod-inoculated calves had a moderate to strong CMI response at 7 dpi (Figure 7C) but went on to develop multiple cutaneous lesions at 11-13 dpi (Figure 6C). This indicates that a strong pre-existing CMI response, as measured by an IGRA, does not always bestow protection against clinical disease.

The IGRA was used on both purified PBMCs and whole blood collected from all 15 calves in group D and groups RS and RA in order to determine if diagnosis of LSD, and particularly diagnosis of preclinical or subclinical LSD, could be improved by using a whole blood IGRA, similar to IFN-γ blood testing of cattle to detect latent tuberculosis (59). However, there were substantial differences in the timing and magnitude of the immune response detected by the whole blood IGRA when compared to the IGRA carried out on PBMCs. This difference was apparent following either needle-inoculation or arthropod-inoculation, with the whole blood IGRA detecting in general fewer positive samples and, in the arthropod inoculated calves, a lower magnitude response. The reason for this difference could be the absence of granulocytes and other blood components such as neutralising antibodies from the purified PBMCs and the stimulation of different immune response pathways by live and SW-UV inactivated LSDV. Overall, the results do not support the development of a whole blood IGRA for LSD diagnosis and, despite the additional time and expense required, this study encourages the use of PBMCs rather than whole blood when carrying out research into the CMI response to LSDV.

The humoral immune response to LSDV has been characterised in numerous experimental studies using both a commercially available ELISA and the virus neutralisation test (VNT). The VNT in these studies often uses tissue culture infectious dose_50_ (TCID_50_) or similar as a readout. In this study we used two new methods which provided more granularity to our assessment of the LSDV humoral immune response – the FVNT, which used number of viral foci rather than TCID_50_ as a read-out, and the B cell ELISpot.

The use of the FVNT to elucidate partial neutralisation provided more in-depth analysis of the humoral immune response in response to needle (n=17) or arthropod (n=10) inoculation. A strong humoral immune response was seen in all clinical calves from 5 dpi onwards with the subsequent increase in neutralising antibodies in the arthropod-challenged calves lagging 6-7 days behind the needle challenged calves, in line with their extended latent period. Both neutralising and binding antibody responses were stronger and more rapid in the clinical calves than the nonclinical calves, regardless of the method of inoculation. This was most obvious at later timepoints (from 15 dpi). A similar pattern of responses has been observed in previous studies (22, 60, 61), and is likely due to the higher virus load in the clinical calves providing more antigenic-stimulation.

This analysis revealed that although the majority of nonclinical calves did not develop antibodies that afforded complete neutralisation by FVNT_100_, there was clear evidence of partially neutralising antibodies in these individual animals from as early as 5 dpi. These partially neutralising antibodies increased gradually over the course of the studies but varied between individual nonclinical animals. Interestingly, two of the nonclinical calves in group D that did not develop completely neutralising antibodies did develop antibodies that neutralised 50 % of LSDV by 21 dpi. A similar pattern was also detected for the two recipient nonclinical calves (RA2 and RS3). This highlights that animals that were negative on the classical VNT_100_ assay that has been used previously for quantifying the humoral immune response to LSDV still have functionally active antibodies present in their sera.

The identification of rapidly appearing partially neutralising antibodies in the two nonclinical arthropod inoculated calves RA2 and RS3 at 7 dpi encouraged us to look more closely at the development of the humoral immune response by developing a B cell ELISpot to monitor the progression of IgM and IgG antibody production. No distinct differences were observed in the IgM and IgG responses from the needle-inoculated animals. However, strong IgM responses were evident in the arthropod-inoculated nonclinical cattle, with a distinct peak at 11 dpi (correlating with the onset of lesions in the clinical animals). This strong IgM response was not observed in the clinical arthropod-inoculated animals. A prominent class switch at 15 dpi was observed in the clinical arthropod-inoculated cattle but occurred later in the nonclinical arthropod-inoculated animals at 23 dpi. These results suggest that early IgM production is a correlate of protection and warrants further analysis. Measuring B-cell antibody responses together with antibody detection determines an earlier immune response and duration of immunity imperative to vaccine surveillance (62).

The role of plasma B cells and antibodies in the primary response to LSDV infection is important for the determination of correlates of protection in infected animals and the development of effective vaccines. Experimental studies investigating the role of neutralising IgG antibodies in mice against vaccinia virus and monkeypox virus identified this antibody isotype developed later in primary infection and after the onset of lesions (63). Similar studies to investigate the role of neutralising IgM in mice challenged with vaccinia virus found that neutralising IgM was detected early in infection and initiated a complement-dependent cascade in the immune response to challenge that could indicate an important role in clinical outcome (63, 64).

This study provides in depth analysis of the adaptive immune response to LSDV through inoculation by intravenous and intradermal inoculation and via the use of arthropod vectors *Stomoxys calcitrans* and *Aedes aegypti* under experimental conditions. The study presents the first evidence of differences in the immune response between clinical and nonclinical cattle, and highlights the importance of using the most relevant model possible when studying disease under experimental conditions. These results will influence the development of improved diagnostic and vaccines for LSDV and for post-vaccination monitoring.

## Author Contributions

PCF, NW, BSB, BC and PMB contributed to the conceptualisation of the study. PCF, NW, HM, BSB, IL, KM, JH, SG and PMB contributed to the study methodology. PCF, NW, IL, KM performed the study validation. PCF, NW, HM and IL performed experimental analysis. PCF, NW, HM, BSB, IL, KM and PMB conducted study investigation. KM provided resource support. BSB and PMB contributed to data curation. PCF, NW, HM, IL and PMB wrote the original draft manuscript and all authors contributed to the final review and editing. IH, KM, AVV, JH, SG and PMB provided supervision and leadership. IH and PMB contributed to project administration. BSB and PMB secured funding and PMB provided the study visualisation.

## Acknowledgements

The authors are grateful to the Animal Services Unit at The Pirbright Institute for their assistance in the husbandry and sampling of cattle over the study period.

