## Supplementary Material for "The immune response to lumpy skin disease virus in cattle is influenced by inoculation route"

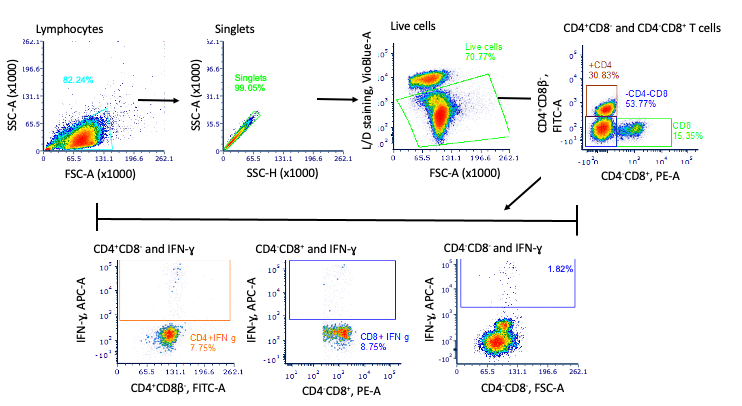

Supplementary Figure 1. Bovine T cell IFN-γ production representative gating strategy for flow cytometric assessment. The flow cytometric gating strategy to assess bovine T cell IFN-γ responses involved assessing the IFN-γ staining of live, singlet, CD4^+^CD8- T cells, CD4-CD8+ T cells and CD4-CD8β- cells. Positive control cells were stimulated with phorbol 12-myristate 13-acetate (PMA) and Ionomycin (Merck). Lymphocytes were surface labelled with LIVE/DEAD™ Fixable Violet dye, anti-bovine CD4-FITC mAb [CC8; Bio-Rad Antibodies], anti-bovine CD8β-PE mAb [CC58; Bio-Rad Antibodies) and intracellularly labelled with an anti-bovine IFN-γ-APC mAb (CC302; Bio-Rad Antibodies) prior to analysis by flow cytometry.

**Supplementary Table 1. Clinical cattle neutralising antibodies following intravenous / intradermal (IV/ID) inoculation.**

|  | Days Post Infection (DPI) | | | | | | | |
| --- | --- | --- | --- | --- | --- | --- | --- | --- |
|  | **0** | **5** | **7** | **9** | **11** | **15** | **17** | **21** |
| Calf ID | **Partial Neutralisation (%)** | | | | | | | |
| A3 | 0 | 34.50 | 36.20 | 88.62 | 92.62 | 100 | 100^*^ |  |
| A5 | 0 | 10.37 | 68.37 | 86.5 | 94.6 | 100 | 100^*^ |  |
| B9 | 0 | 16.75 | 22.75 | 45.62 | 72.75 | 100 | 100 | 100 |
| C12 | 0 | 16.13 | 15.38 | 38.00 | 40.00 | 90.75 | 100 | 100 |
| D2 | 0 | 20.00 | 34.37 | 56.12 | 85.50 | 100 | 100 | 100 |
| D4 | 0 | 21.50 | 18.12 | 77.75 | 91.25 | 100 | 100 | 100 |
| D5 | 0 | 14.50 | 37.25 | 72.00 | 93.5 | 100 | 100 | 100 |
|  | **Complete Neutralisation Titres (FVNT_100_)** | | | | | | | |
| A3 | 0 | 0 | 0 | 10 | 60 | 160 | 480^*^ |  |
| A5 | 0 | 0 | 15 | 30 | 60 | 80 | 120^*^ |  |
| B9 | 0 | 0 | 0 | 0 | 15 | 80 | 640 | 960 |
| C12 | 0 | 0 | 0 | 0 | 0 | 30 | 240 | 640 |
| D2 | 0 | 0 | 0 | 0 | 0 | 120 | 320 | 960 |
| D4 | 0 | 0 | 0 | 0 | 20 | 80 | 960 | 1920 |
| D5 | 0 | 0 | 0 | 0 | 15 | 80 | 480 | 2560 |

^*^ Human endpoint

**Supplementary Table 2. Nonclinical cattle neutralising antibodies following intravenous / intradermal (IV/ID) inoculation.**

|  | Days Post Infection (DPI) | | | | | | | |
| --- | --- | --- | --- | --- | --- | --- | --- | --- |
|  | **0** | **5** | **7** | **9** | **11** | **15** | **17** | **21** |
| Calf ID | **Partial Neutralisation (%)** | | | | | | | |
| A1 | 0 | 0 | 0 | 0 | 0 | 0 | 0 | 0 |
| A2 | 0 | 16.20 | 17.10 | 70.25 | 88.25 | 51.00 | 63.60 | 100 |
| A4 | 0 | 21.37 | 28.60 | 84.25 | 88.00 | 72.25 | 95.00 | 100 |
| B6 | 0 | 0 | 0 | 0 | 0 | 0 | 0 | 0 |
| B7 | 0 | 0 | 10.75 | 40.75 | 39.62 | 62.25 | 88.12 | 100 |
| B8 | 0 | 9.25 | 21.25 | 37.62 | 40.62 | 80.0 | 88.00 | 100 |
| B10 | 0 | 0 | 22.65 | 11.50 | 47.00 | 72.25 | 87.87 | 100 |
| C11 | 0 | 0 | 0 | 0 | 0 | 0 | 0 | 0 |
| C13 | 0 | 0 | 8.63 | 14.75 | 16.00 | 69.62 | 70.00 | 100 |
| C14 | 0 | 0 | 20.88 | 22.63 | 24.00 | 51.13 | 52.00 | 54.38 |
| C15 | 0 | 50.88 | 58.00 | 67.38 | 68.00 | 70.10 | 75.00 | 92.88 |
| D1 | 0 | 48.75 | 46.00 | 63.12 | 84.75 | 94.37 | 100 | 100 |
| D3 | 0 | 28.12 | 22.75 | 59.62 | 87.87 | 96.87 | 100 | 100 |
|  | **Complete Neutralisation Titres (FVNT_100_)** | | | | | | | |
| A1 | 0 | 0 | 0 | 0 | 0 | 0 | 0 | 0 |
| A2 | 0 | 0 | 0 | 0 | 0 | 10 | 10 | 10 |
| A4 | 0 | 0 | 0 | 0 | 0 | 15 | 40 | 120 |
| B6 | 0 | 0 | 0 | 0 | 0 | 0 | 0 | 0 |
| B7 | 0 | 0 | 0 | 0 | 0 | 15 | 20 | 40 |
| B8 | 0 | 0 | 0 | 0 | 0 | 15 | 30 | 80 |
| B10 | 0 | 0 | 0 | 0 | 0 | 10 | 20 | 20 |
| C11 | 0 | 0 | 0 | 0 | 0 | 0 | 0 | 0 |
| C13 | 0 | 0 | 0 | 0 | 0 | 15 | 15 | 15 |
| C14 | 0 | 0 | 0 | 0 | 0 | 0 | 0 | 0 |
| C15 | 0 | 0 | 0 | 0 | 0 | 10 | 10 | 10 |
| D1 | 0 | 0 | 0 | 0 | 0 | 160 | 60 | 160 |
| D3 | 0 | 0 | 0 | 0 | 0 | 60 | 160 | 640 |

**Supplementary Table 3. Recipient clinical cattle neutralising antibodies following arthropod inoculation**

|  | Days Post Infection (DPI) | | | | | | | | | | |
| --- | --- | --- | --- | --- | --- | --- | --- | --- | --- | --- | --- |
|  | **0** | **5** | **7** | **9** | **11** | **15** | **17** | **21** | **23** | **25** | **29** |
| Calf ID | **Partial Neutralisation (%)** | | | | | | | | | | |
| RA1 | 0 | 20.25 | 18.62 | 30.37 | 51.50 | 89.92 | 93.57 | 100^*^ |  |  |  |
| RA3 | 0 | 0 | 16.87 | 0 | 24.87 | 78.87 | 91.87 | 96.12 | 100^*^ |  |  |
| RA4 | 0 | 0 | 15.87 | 0 | 36.87 | 79.50 | 87.75 | 91.37 | 100 | 100 | 100 |
| RA5 | 0 | 0 | 0 | 11.25 | 54.12 | 76.75 | 90.00 | 96.62 | 100 | 100^*^ |  |
| RS1 | 0 | 0 | 14.25 | 0 | 68.00 | 86.00 | 94.50 | 100 | 100 | 100^*^ |  |
| RS2 | 0 | 0 | 11.62 | 7.87 | 63.75 | 86.00 | 92.75 | 100 | 100 | 100^*^ |  |
| RS4 | 0 | 11.00 | 0 | 0 | 59.50 | 89.00 | 94.12 | 100^*^ |  |  |  |
| RS5 | 0 | 0 | 27.25 | 0 | 66.75 | 71.00 | 75.00 | 64.20 | 88.20 | 94.50 | 63.12 |
|  | **Complete Neutralisation Titres (FVNT_100_)** | | | | | | | | | | |
| RA1 | 0 | 0 | 0 | 0 | 0 | 20 | 120 | 320^*^ |  |  |  |
| RA3 | 0 | 0 | 0 | 0 | 0 | 0 | 20 | 20 | 480^*^ |  |  |
| RA4 | 0 | 0 | 0 | 0 | 0 | 0 | 10 | 80 | 640 | 1280 | 1280 |
| RA5 | 0 | 0 | 0 | 0 | 0 | 0 | 15 | 120 | 1280 | 1280^*^ |  |
| RS1 | 0 | 0 | 0 | 0 | 0 | 0 | 40 | 480 | 1280 | 960^*^ |  |
| RS2 | 0 | 0 | 0 | 0 | 0 | 0 | 30 | 240 | 1280 | 1280^*^ |  |
| RS4 | 0 | 0 | 0 | 0 | 0 | 0 | 80 | 1280^*^ |  |  |  |
| RS5 | 0 | 0 | 0 | 0 | 0 | 0 | 0 | 0 | 10 | 15 | 0 |

^*^ Human endpoint

**Supplementary Table 4. Recipient nonclinical cattle neutralising antibodies following arthropod inoculation**

|  | Days Post Infection (DPI) | | | | | | | | | | |
| --- | --- | --- | --- | --- | --- | --- | --- | --- | --- | --- | --- |
|  | **0** | **5** | **7** | **9** | **11** | **15** | **17** | **21** | **23** | **25** | **29** |
| Calf ID | **Partial Neutralisation (%)** | | | | | | | | | | |
| RA2 | 0 | 28.00 | 44.50 | 25.62 | 36.62 | 62.12 | 63.25 | 55.37 | 52.50 | 54.90 | 51.60 |
| RS3 | 0 | 19.87 | 28.87 | 18.12 | 68.37 | 74.25 | 79.37 | 64.25 | 72.30 | 81.50 | 40.37 |
|  | **Complete Neutralisation Titres (FVNT_100_)** | | | | | | | | | | |
| RA2 | 0 | 0 | 0 | 0 | 0 | 0 | 0 | 0 | 0 | 0 | 0 |
| RS3 | 0 | 0 | 0 | 0 | 0 | 0 | 0 | 0 | 0 | 0 | 0 |
